# Comparative mutant analyses reveal a novel mechanism of ARF regulation in land plants

**DOI:** 10.1101/2023.11.09.566459

**Authors:** Michael J. Prigge, Nicholas Morffy, Amber de Neve, Whitnie Szutu, María Jazmín Abraham-Juárez, Kjel Johnson, Nicole Do, Meirav Lavy, Sarah Hake, Lucia Strader, Mark Estelle, Annis E. Richardson

## Abstract

A major challenge in plant biology is to understand how the plant hormone auxin regulates diverse transcriptional responses throughout development, in different environments, and in different species. The answer may lie in the specific complement of auxin signaling components in each cell. The balance between activators (class-A AUXIN RESPONSE FACTORS) and repressors (class-B ARFs) is particularly important. It is unclear how this balance is achieved. Through comparative analysis of novel, dominant mutants in maize and the moss *Physcomitrium patens*, we have discovered a ∼500-million-year-old mechanism of class-B ARF protein level regulation, important in determining cell fate decisions across land plants. Thus, our results add a key piece to the puzzle of how auxin regulates plant development.

## Main Text

Plants develop post-embryonically through the activity of stem cells. Pattern formation, differentiation, and tissue deformation define the shape of complex organs, while iterative development of these organs build the plant body. Modulation of these developmental processes gives rise to a huge diversity in organ shapes and plant architectures. Despite this complexity, a single phytohormone, auxin, has a major, conserved role across land plants (*1–4*), underpinning cell fate decisions, organ initiation, shape and plant architecture. How auxin is able to precisely regulate so many distinct processes is one of the biggest questions in plant biology.

Canonical auxin signaling occurs through the activity of a suite of deeply conserved transcriptional regulators found in all land plants (*3*, *5*, *6*). The AUXIN/INDOLE-3-ACETIC-ACID repressors (AUX/IAAs), AUXIN RESPONSE FACTORS (ARFs) and the TRANSPORT INHIBITOR RESPONSE 1 /AUXIN-SIGNALING F-BOX (TIR1/AFB) co-receptors are components of the best-characterized pathway of auxin-mediated transcriptional regulation (reviewed by (*7–9*)). All three components are in multigene families that have greatly expanded in flowering plants (*3*). When auxin levels are low, AUX/IAA proteins are bound to ARFs, and recruit transcriptional co-repressors preventing ARF activation of auxin responsive genes. In the presence of elevated levels of auxin, the TIR1 co-receptor forms a complex with auxin and AUX/IAA repressors, resulting in ubiquitination and degradation of AUX/IAAs via the 26S proteasome (*10–12*). Loss of the AUX/IAAs relieves ARF repression and facilitates auxin transcriptional responses (*13*). Land plant ARFs are divided into three phylogenetic lineages: class-A, -B and -C. Each class is associated with particular functions based on protoplast studies: class-A ARFs are activators, while class-B and -C are potential repressors (*14*, *15*). All ARFs share a DNA binding domain (DBD), a middle domain and a C-terminal Phox and Bem1 (PB1) domain (*16*). Differences in domain sequences lead to variation in DNA binding activity (*17–21*), interacting partners (including other ARFs, the AUX/IAAs, other transcription factors, and chromatin remodelers (*16*)), and the ability to oligomerize (*22*). Modulation of auxin response is believed to be determined by the complement of ARFs and AUX/IAAs in the cell, and *in vitro* analysis has revealed different activation levels of ARF–AUX/IAA combinations indicating potential mechanisms for sub-functionalization (*23*). However, the elegant model of ARF de-repression in the presence of auxin is not able to account for all auxin responses; only 5 of the 22 Arabidopsis ARFs have been shown to functionally interact with AUX/IAAs *in vivo* (*16*). Alternative methods of ARF regulation are required for the diverse auxin responses that are crucial for plant development, particularly in relation to the class-B ARFs that do not fit this model that is largely based on class-A ARF activity.

Although genetic studies implicate class-B function in diverse developmental processes (*24–26*), their mode of action and regulation is unclear. Studies in non-vascular species like Marchantia and Physcomitrium indicate that class-B ARFs can directly compete with class-A to bind to promoter sequences (*12*, *27*). Genome-wide analyses in flowering plants support the hypothesis that competition between class-A and -B ARFs may be important for the regulation of a significant fraction of auxin-regulated genes across land plants. (*20*, *28*, *29*). In addition, class-B ARFs can function independently of AUX/IAAs directly recruiting the TOPLESS (TPL) repressor to chromatin (*30–34*), providing an AUX/IAA-independent way to repress gene expression. Regardless of mechanism, the relative level of class-A and -B ARFs is a key determinant of auxin regulated transcription. Without a clear understanding of class-B ARF regulation, and how it has evolved, we lack a key piece of the auxin regulatory system in plants.

To address this deficiency, we leveraged a comparative approach by investigating dominant class-B ARF mutants in the distantly related species *Zea mays* (maize) and the moss *Physcomitrium patens* (*P.patens*) which diverged ∼500 mya (*35*). The mutations result in amino acid substitutions in a loop region located in the DBD of these proteins. We demonstrate that this loop region contains a novel degron that is required for regulation of class-B ARF accumulation thus, revealing a new, deeply conserved mechanism of class-B ARF regulation in land plants.

### *Truffula* has pleiotropic defects

Maize is a classic model genetic system, with a rich history of developmental mutant analyses (*36*). We found the striking, dominant *Truffula* (*Trf*) mutant in an EMS-mutagenized population and analyzed it in two inbred backgrounds, Mo17 and W22. In both inbreds, when compared to normal siblings, *Trf* mutants have reduced plant height, increased leaf number, and feminization of the male tassel (Fig.1A-D, fig.S1). The mutation did not transmit through the female in Mo17, but did at a reduced rate in W22, permitting analysis of homozygotes in W22. Mutant plants are half the height of normal siblings in W22 (Fig.1A) and two-thirds the size in Mo17 (Fig.1B). The reduction in height results from shorter internodes, the stem section between leaves (fig.S1B-C). In Mo17, non-mutant siblings produce 15 leaves while heterozygous mutants produce an average of 19. In W22, non-mutants produce 15 leaves, heterozygotes produce 29 and homozygous mutants produce an average of 54 (Fig.1E). Analysis of flowering time revealed that mutants flower at the same time as normal siblings, and therefore the increase in leaf number is due to a shorter plastochron in *Trf* (the time between leaf initiations) (fig.S1A). Additional *Trf* phenotypes were inbred dependent such as defects in midrib formation (15% of leaves show midrib defects) ranging from midribless leaves to multiple midribs (Fig.1C, arrowheads) and in leaf shape, forming very rare tube leaves in W22 (<1% of leaves). The feminization in W22 was a few kernels at the base of the male tassel while in Mo17 the feminization was extreme with ears wrapped in husk leaves replacing the tassel branches (Fig.1D).

**Fig. 1.**
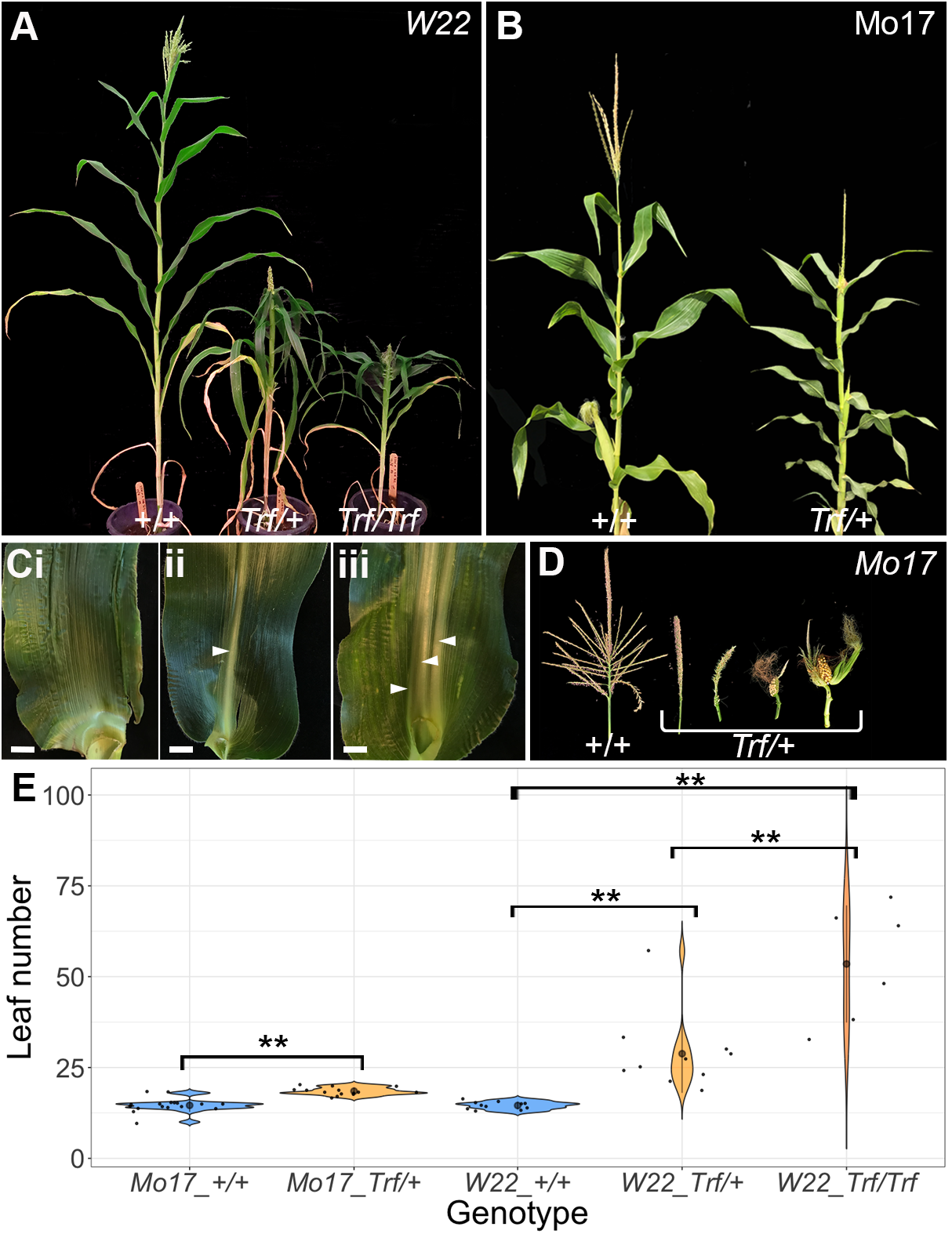
Maize *Truffula* mutants have pleiotropic defects. Normal (*+/+*) *and Trf* siblings (*Trf/+*, *Trf/Trf*) in the W22 (**A,C**) and Mo17 (**B,D**) inbred backgrounds. (**A-B**) Mature plants. (C) Midrib phenotypes in *Trf* leaves: midribless (i), normal (ii), and multiple midribs (iii). (D) Tassel phenotypes showing feminisation in *Trf* siblings. (**E**) Leaf number in mature plants of normal (blue) and *Trf* siblings (orange) (“**” p-value <0.05, n≥6, black dot and bars: mean ± sd.)

### *Trf* is associated with a mutation in *ZmARF28*

To identify the underlying causal mutation in *Trf* we carried out whole genome sequencing bulk segregant analysis (*37*), identifying a region between 11.0 Mbp and 20.0 Mbp on chromosome 10 (Fig.2A). Short sequence repeat (SSR) mapping (Fig.2B, Table S1) refined the region to between 13.5Mbp and 14.0Mbp. This fine-mapping region contains 8 annotated genes, of which 5 are expressed in the vegetative shoot apex according to RNAseq (Fig.2B, fig.S2A, Table S2). Analysis of the WGS data in the narrow mapping region reveals 188 EMS-type variants (G:C to A:T transitions) specific to *Trf* siblings. SNPEff (*38*) analysis predicts 7 moderate effect EMS-type SNPs in the interval (Table S3). Of these, only a single SNP in exon 8 of *ZmAUXIN RESPONSE FACTOR28 (ZmARF28*, Zm00001eb408800), resulting in a Ser-to-Asn change in the DBD (Fig.2C- D), is *Trf* specific when compared to other maize inbred backgrounds (Table S3). Thus, suggesting that this mutation in *ZmARF28* is responsible for the *Trf* phenotype.

**Figure 2:**
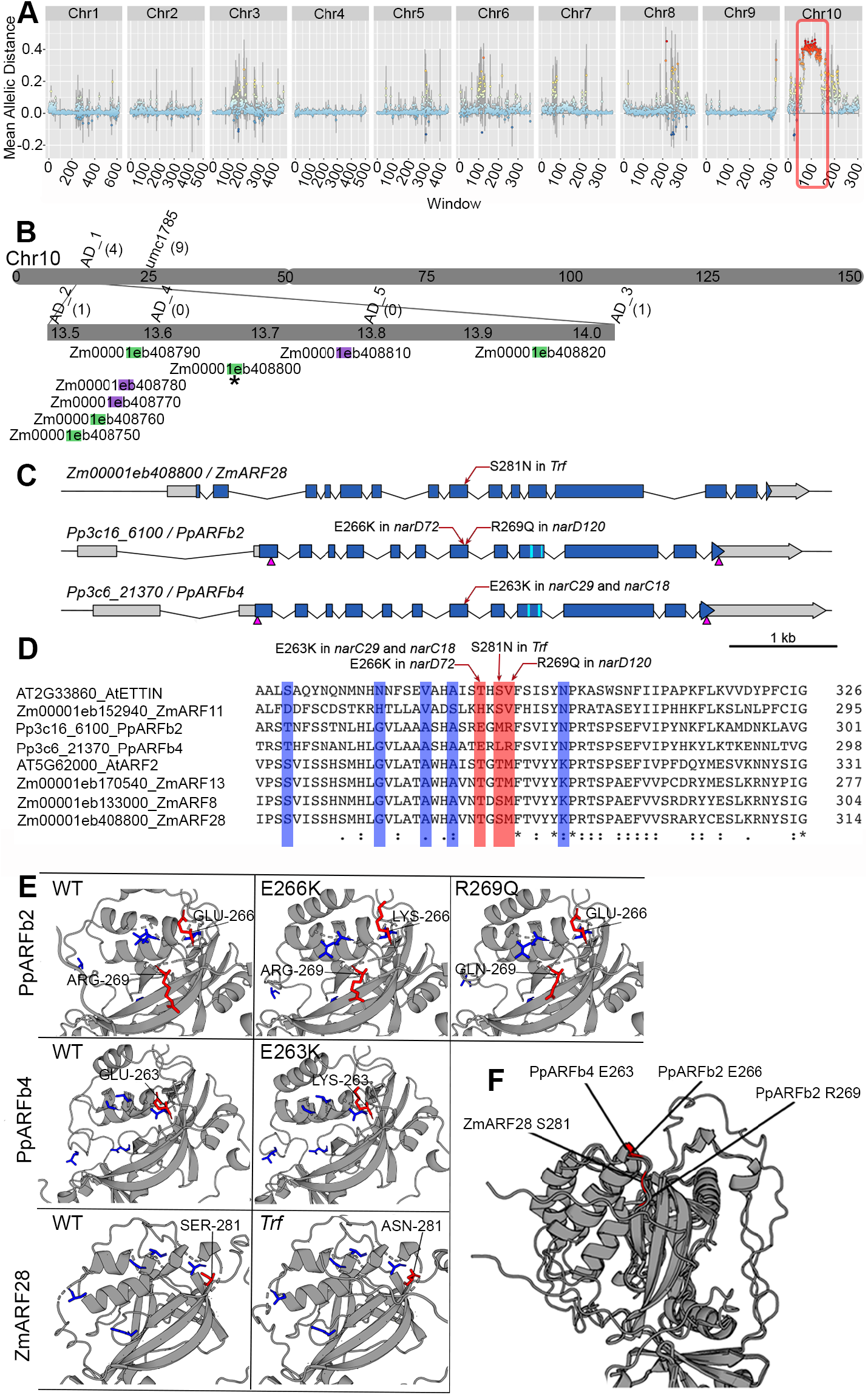
T**h**e ***Truffula* lesion maps to ZmARF28.** (**A**)Plotting WGS-BSA variant mean allelic distances in 0.5Mbp windows across the maize NAM5.0 reference genome, reveals a mapping location on chromosome 10 which is more associated with *Trf* mutants than normal siblings (red box).(**B**) PCR mapping narrows the mapping region to between 13.5Mbp and 14.0Mbp on chromosome 10. Mapping primer positions are indicated above (AD_# and umc#), and recombinant numbers are shown in brackets. There are 8 genes in the interval, of which 5 have an FPKM >1 in our shoot apical meristem RNAseq dataset (green). A moderate effect, *Trf*-specific, EMS-type SNP is present in Zm00001eb408800 (ZmARF28). (**C**)The locations of the *Trf, narD72, narD120, narC29*, and *narC18* mutations (red arrows). The exons: blue, 5’ and 3’UTRs (grey), CRISPR sgRNA targets for deletions (magenta arrowheads), and miRNA regulatory sites (cyan) are indicated. (**D-F**) The maize *Trf* mutation, and the moss *narD72, narD120, narC29*, and *narC18* mutations map to the same loop-region in the ARF protein. (**D**) Multiple sequence alignments of ARFs from maize, Arabidopsis, and moss. Mutation locations (red), and amino acids previously shown to be important for DNA binding (blue) are indicated. (**E-F**) Homology modelling of ZmARF28 DNA binding domain for both wild-type and *Trf* mutant protein.

To investigate further, we carried out an RNAseq analysis. Consistent with a defect in auxin regulated transcriptional response, RNAseq analysis of vegetative shoot apices found 1804 significantly differentially expressed genes (sigDE, padj<0.05, and log2 fold change <-05 or >0.5) with enrichment for GO terms associated with auxin signaling and response, and transcriptional control. In line with the phenotype, we also observed enrichments for terms associated with meristem development (fig.S2C). Crucially, this RNAseq analysis showed that ZmARF28 is not significantly differentially expressed in *Trf* vs normal siblings (fig.S2B), suggesting that the *Trf* mutation affects protein activity or post-translational regulation of ZmARF28.

As ZmARF28 is a co-orthologue of Arabidopsis AtARF2 (fig.S5), we carried out sequence alignments (Fig.2D) and homology modelling. This analysis indicates that the *Trf* mutation is outside of the core residues required for DNA binding (Fig.2D-E, blue, (*17*)) in an externally facing loop region (Fig.2E-F). These findings suggest that the novel *Trf* phenotype is not due to a mutation in an already characterized ARF motif or domain, potentially revealing a novel regulatory module.

Although protein structure is predicted to be conserved, amino acid sequence is not highly conserved in this region of plant class-B ARFs (Fig.2D). This suggests that the *Trf* phenotype could be due to the disruption of a grass-specific regulatory pathway in the expanded family of class-B ARFs. Alternatively, the *Trf* mutation could highlight the role of a functionally conserved region of class-B ARFs that has yet to be described due to lack of sequence conservation.

### Mutations in the same domain of class-B ARFs in *P. patens* cause auxin resistance

To determine whether the *Trf* mutation revealed a grass-specific class-B ARF regulatory pathway, or a core part of the auxin module that evolved early in land plant evolution, we looked for independent class-B ARF mutants in the moss *Physcomitrium patens*. There are four class-B *ARF* genes in this species, *PpARFb1* to *PpARFb4.* We found that the genomes of four newly isolated *P.patens NAA-RESISTANT (nar* (*39*)) mutants had missense mutations in the same *Trf* region in two ARFs (*PpARFb2* & *PpARFb4*, Fig.2C- F). *narC29* and *narC18* mutants have missense mutations causing Glu-263 to lysine substitutions (E263K) in PpARFb4/Pp3c6_21370, while *narD72* has a substitution of the corresponding Glu-266 of PpARFb2/Pp3c16_6100 (E266K). *narD120* has a glutamine substitution at Arg-269 of PpARFb2 (R269Q). When the DBD are aligned, the residues affected in the moss mutants are proximal to the position altered by the maize *Trf* mutation (Fig.2D,F). These class-B ARF moss *nar* mutants had clear defects in the auxin-regulated transition from highly photosynthetic chloronemal cells to “nutrient foraging” spreading caulonemal cells and a subsequent delay in the production of 3D-leafy shoot (gametophore) growth forms. Whereas wild-type produce ectopic reddish brown rhizoid cells on NAA-containing medium, the *nar* mutants produce gametophores even when grown on media containing micromolar amounts of NAA (Fig.3A). Further supporting the reduction in auxin response in these *nar* mutants, expression of the *proDR5:DsRed2* auxin-signaling reporter was also dramatically reduced in the *narC29*, *narC18, narD72* mutants, and to a lesser extent in the *narD120* mutant (Fig.3A).

**Figure 3:**
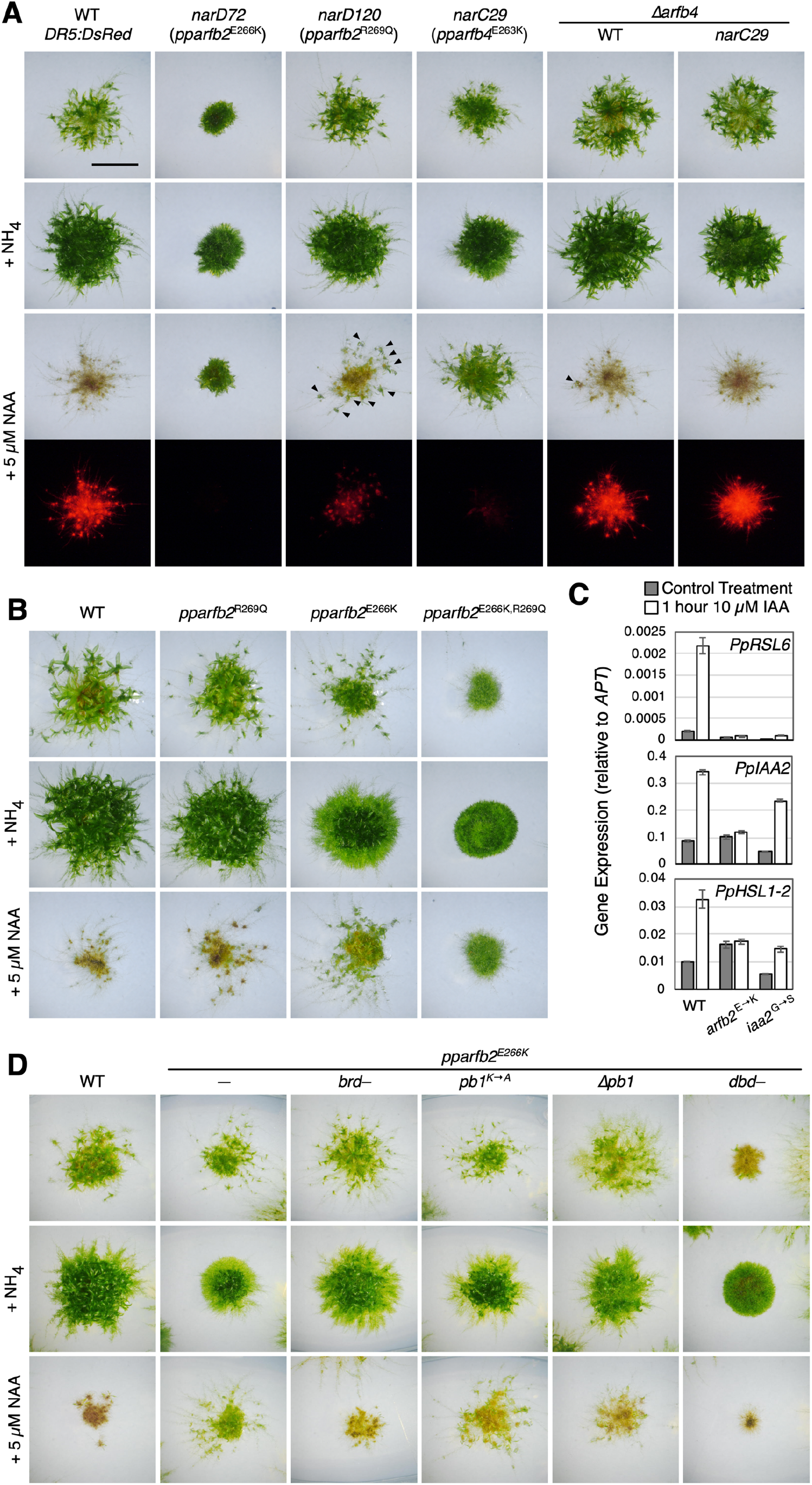
M**u**tations **in *PpARFb2* and *PpARFb4* cause auxin resistance.** (**A**) Comparison of wild type, *nar* mutants, and *Δpparfb4* deletion mutants after 21 days of growth on standard medium (BCD) and BCD supplemented with 5 mM ammonium tartrate or 5 µM NAA. The plants grown on NAA were also imaged for RFP fluorescence from the *proDR5:DsRed* auxin-response reporter. On NAA-containing media, wild-type and *Δpparfb4* strains produce ectopic rhizoids and high levels of RFP fluorescence whereas the *nar* mutants produce green leafy gametophores. Arrowheads indicate leafy shoots in *narD120* and *Δpparfb4*. Scale bar is 5 mm. (**B**) Comparison of wild type and gene-edited point mutants grown for 21 days, as in panel (A). (**C**) Expression of select auxin-responsive genes (*PpRSL6, ROOT HAIR DEFECTIVE SIX-LIKE 6*; *PpIAA2, Aux/IAA 2*; *PpHSL1-2*, *HOOKLESS1*-*LIKE 2*) in wild type, *pparfb2^E266K^,* and auxin-response mutant *ppiaa2^G325S^* grown for 1h on media containing 10 µM IAA or 0.01% ethanol. Transcript levels were normalized to the *APT* gene. (**D**) Growth comparisons of wild type, the *pparfb2^E266K^* mutant, and the *pparfb2^E266K^* mutant with second-site mutations, as in panel (A). *brd–*, L614A+F615A substitutions in the B3 Repression Domain; *pb1K→A*, K682A substitution in the PB1 domain; *Δpb1*, a 5 bp frameshift-causing deletion starting at G680 codon; *dbd–*, P194A+R196A substitutions affecting DNA-interacting residues.

To independently verify that *pparfb* mutations are responsible for the above *nar* phenotypes, we used oligo-mediated gene editing (*40*) to introduce the E-to-K substitution to all four *PpARFb* loci. The *pparfb2^E^*^266^*^K^*and *pparfb4^E^*^263^*^K^*lines recapitulated the *nar* phenotypes of *narD72* and *narC29*, respectively, and the *pparfb1^E266K^* and *pparfb3^E266K^*lines were also significantly resistant to NAA (Fig.3B; fig.S3B). *pparfb2^R269Q^*and *pparfb4^R266Q^* mutations conferred weak *nar* phenotypes like *narD120* (Fig. 3B; fig.S3C). Interestingly, combining both the E-to-K and R-to-Q mutations in the same gene resulted in dramatically stronger phenotypes approaching those of quadruple mutants lacking all four *TIR1/AFB* genes which are completely auxin insensitive (Fig.3B; fig.S3C). In-frame deletions and insertions in this region of *PpARFb2* also conferred either intermediate phenotypes similar to *pparfb2*^E266K^ or strong phenotypes similar to *pparfb2*^E266K+R269Q^ (Table S4). Reduced expression of *proDR5:DsRed* and reduced auxin induction of auxin regulated genes in *pparfb2*^E266K^ confirmed that the phenotypes are due to reduced auxin response (Fig.3C, fig.S3D–E).

The phenotypes of the four *nar* mutants are similar to lines expressing *PpARFb* genes with mutated *miR1219* and *tasiARF* targets which accumulate PpARFb (*41*). The *nar* mutations do not affect any known small RNA regulatory elements, but the phenotypic similarities suggest that the *nar* lesions represent dominant mutations. To test this, we used CRISPR/Cas9 to delete the mutated genes in the mutant lines. Deletion of either of the wild-type *PpARFb2* and *PpARFb4* genes in moss has very minor effects on phenotype (*12*) but deletion of the mutated genes in the *nar* mutants restored sensitivity to NAA indicating that, like the maize *Trf* mutant, the *P.patens arfb2* and *arfb4* variants are dominant mutations (Fig.3A; fig.S3A) with defects in auxin signaling.

### All major domains are required for the dominant phenotype

To understand how the mutations confer gain-of-function phenotypes in moss, we introduced second-site mutations affecting key residues of known functional domains into the *pparfb2^E266K^*mutant (Fig.3D; fig.S3E–F). Mutating the BRD (L614S+F615S) eliminates the interaction with a TPL homolog (TPL2/Pp3c9_21250) and reverts the *pparfb2^E266K^* phenotype to wild-type (Fig.3D; fig.S3F–G). Similarly, mutating a conserved lysine residue in the positive face of the PB1 domain (K682A) partially suppresses the *arfb2^E266K^*phenotype, and a frame-shift mutation near the start of the domain results in near-complete suppression. Introducing a pair of substitutions (P194A+R196A) shown to abolish DNA binding (*17*), restores NAA response, although additional phenotypes appear suggesting dominant-negative effects. None of these functional-domain mutations caused phenotypic effects in an otherwise wild-type *PpARFb2* (fig.S3F–G). This shows that the gain-of-function phenotypes depend on all known ARF protein functions; DNA binding, TPL recruitment, and PB1-mediated oligomerization.

### Mutations increase class-B ARF stability

Based on the phenotypes, and lack of *ZmARF28* and *PpARFb* expression changes in the mutants, we hypothesized that the maize *Trf* and *P.patens arfb* mutant phenotypes were due to accumulation of ZmARF28 and PpARFb protein respectively.

To test whether the PpARFb proteins are more abundant in the moss mutants we knocked-in mYPet yellow-fluorescent protein tags to the four endogenous *PpARFb* loci and the four E-to-K mutants allowing us to visualize and quantify protein abundance. The YFP signal dramatically increases in all nuclei of the E266K/E263K mutants compared to the weak and sporadic YFP signal detected in nuclei in the *mYPet*-tagged wild-type loci (Fig.4A-D). Since the transcript levels are similar in the mutant and wild-type lines (fig.S4A) this increased signal suggests that the mutations affect PpARFb stability. To determine whether the wild-type proteins are subject to degradation via the 26S proteasome, we treated the wild-type mYPet lines overnight with the proteasome inhibitor, Bortezomib. Quantification of the mYPet signal revealed significant increases in protein level for all four wild-type lines in the presence of the inhibitor (Fig.4C-D). These results indicate that the ARFb proteins are subject to proteasome-dependent degradation and that this degradation is disrupted by the dominant mutations.

**Figure 4:**
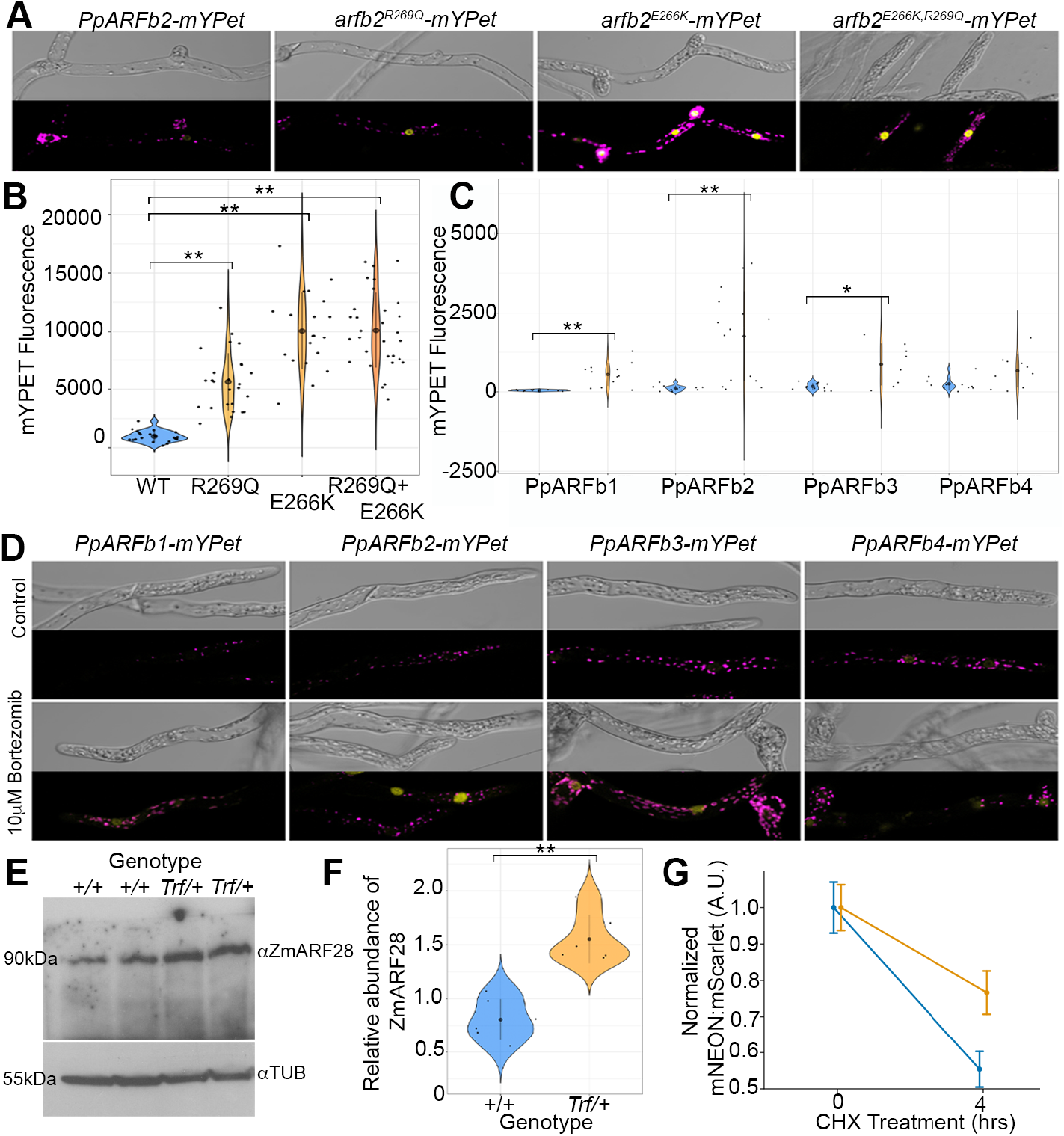
M**u**tations **in the same domain result in stabilization of ZmARF28, PpARFb, and AtARF2.** (**A-B**) PpARFb2-mYPet signal in *P.patens*, in wildtype, pparfb2^R269Q^, pparfb2^E266K^, and the pparfb2^E226K,R269Q^ double mutants. (**A**) Confocal microscopy images (PpARFb2-mYPet, yellow; chlorophyll autofluorescence, magenta), and quantification (**B**). (**C-D**) Class-B ARF accumulation in *P.patens* in control conditions (blue) compared to treated with 10 µM Bortezomib (orange). (**D**) Confocal microscopy images (PpARFb-mYPet, yellow), and quantification (**C**). (**E**) Western blot of ZmARF28 in *+/+* and *Trf/+* mutant siblings from a W22 backcross population. Lanes are different biological replicates. antiTubulin is used as a loading control. (**F**) Quantification of ZmARF28 abundance relative to TUBULIN. (“**” p-value <0.01, “*” p-value<0.05, black dot and bar: mean ± sd.) (**G**) Mean (±SEM) ratiometric signal of AtARF2- mNeonGreen (blue) and AtARF2-Trf-mNeonGreen (orange) in Arabidopsis protoplasts, relative to mScarlet after cyclohexamide (CHX) treatment.

To determine if protein levels were also elevated in the maize *Trf* mutants, we developed an antibody against ZmARF28. Semi-quantitative Western blot analysis of whole-protein extract from vegetative seedlings revealed a statistically significant increase in abundance of ZmARF28 protein in the *Trf* mutant compared to normal siblings (Fig.4E- F), supporting the hypothesis that ZmARF28 is stabilized in *Trf*. This suggests that both the *Trf* and moss mutants disrupt a deeply conserved degron motif.

To further test the effect of the *Trf* mutation on protein stability in flowering plants, we expressed the Arabidopsis ortholog of ZmARF28, AtARF2, and a *Trf* variant of AtARF2 (arf2^T298N^) in Arabidopsis mesophyll protoplasts using a ratiometric reporter system. Protoplasts expressing AtARF2 or arf2^T298N^ were treated with the transcriptional inhibitor cycloheximide (CHX). Similar to AtARF1 (*42*), AtARF2 shows a half-life of approximately 4-hours. Conversely, arf2^T298N^ only lost 23% of its initial protein level after 4-hours of CHX treatment, suggesting that its half-life is longer than wild-type AtARF2 (Fig.4G). These results show that the *Trf* mutation in AtARF2 also increases overall protein stability, further supporting functional conservation of this region.

## Discussion

The precise complement and abundance of auxin signaling components in each cell determines auxin sensitivity and how the auxin signal is decoded to produce distinct transcriptional responses. Data across land plants shows that the relative levels of class-A and -B ARFs have a key role in regulating auxin response (*4*, *12*). We leveraged a comparative analysis of species that diverged ∼500mya and identified an ancient regulatory mechanism that controls class-B ARF accumulation. By combining our findings with an emerging understanding of differential class-A and -B binding preferences (*20*, *28*, *29*), we can start to build a model of how class-A and -B ARF function may be regulated.

It is clear from our comparative mutational analysis, that the ratio of class-A to -B ARFs is essential in dictating cell fate decisions in land plants. Elevated class-B ARF, as in the dominant *Pparfb* mutations, pushes the apical cells towards a leafy gametophore fate, despite nutrient deficiency. Similarly in maize, class-B ARF ZmARF28 level influences meristem cell fate, leaf medio-lateral patterning, and sex determination. Defects in these auxin-determined decisions lead to changes in phyllotaxy, node elongation and inflorescence architecture whilst fate decisions like the transition to flowering remain unaffected, highlighting the diversification of ARF function in flowering plants.

It is remarkable that, despite a lack of sequence-level conservation, the same short region within the DBD of class-B ARFs has an essential role in regulating protein level across land plants. Assuming that this regulatory region interacts with a second protein, perhaps a ubiquitin protein ligase, it is possible that the degron sequence and interacting protein have co-evolved during the ∼500my that separate maize and *P.patens,* explaining differences in sequence and response. This further emphasizes that, like class-B *ARFs* regulation by the *TAS3* tasiRNA, some aspects of ARF regulation are ancient, although the outcomes can be different; in flowering plants, *TAS3* regulation of *ARF3* and *ARF4* is required for adaxial-abaxial patterning, while in *P.patens* the pathway regulates protonemal development (*41*, *43*, *44*). These differences further highlight lineage-specific changes to auxin-regulated cell fate decisions.

Our work underscores the importance of protein-level regulation in tuning auxin response. Previous work found that proteasome-regulation of class-B AtARF1 and AtARF2 stability in Arabidopsis, is dependent on AtHOOKLESS1 activity (*45*) unlike our results here. More recently, the F-box protein AtAFF1 was shown to regulate the levels and behavior of the class-A ARFs, AtARF7 and AtARF19 (*46*). Further instances of proteasome regulation of ARFs, potentially conserved across evolution as in the case of the *Trf* and *nar* mutants are likely. It will be particularly interesting to see if similar non-sequence conserved regulatory mechanisms are common in the ARF lineages.

The TIR1/AFB-Aux/IAA pathway is subject to complex regulation consistent with its essential role in diverse processes. These parallel regulatory pathways enhance auxin response robustness, and modulation likely facilitated the expansion and fine-tuning of auxin-mediated processes across land plant lineages. Comparative work, like ours, is key to the discovery of the auxin puzzle pieces and how they fit together.

## Acknowledgments

We thank Gerry Neuffer for the kind donation of the original *Trf* seed. We thank Samuel Leiboff (Oregon State University), Madelaine Bartlett (UMass Amherst), Dolf Weijers (Wageningen University), Sandy Hetherington (University of Edinburgh) for their discussions and comments. We also thank the ARS/USDA Plant Gene Expression Center plant growth support staff, the University of Edinburgh Plant Growth Facility, and CYVERSE (work supported by the National Science Foundation under Award Numbers DBI-0735191, DBI-1265383, and DBI-1743442. URL: www.cyverse.org) for technical support and resources.

## Funding

University of Edinburgh Start-Up Fund (AER) UKRI EP/Y010116/1 (AER)

National Science Foundation IOS1922543 (SH), PGRP BIO-2112056 (LCS)

National Institutes of Health R01 GM043644 (ME), R35GM141892 (ME), R35 GM136338 (LCS)

Ledell Family Research Scholarship for Science & Engineering (WS)

## Author contributions

Conceptualization: AER, LS, ME, SH, NM, MP Methodology: HP, FTGS, CW, JRK, NJB, PRB, JLS, EH

Investigation: SH, AN, KJ, AER cloned and phenotyped *Trf*, AER carried out the WGS and RNAseq analyses. NM carried out the protoplast stability experiments and the AlphaFold analysis. MAJ purified the ZmARF28 antibody and carried out the western blots. WS carried out the nar mutant screen and the initial mutant characterization with help from MJP. MJP generated and characterized all moss deletion, genome-edited, and reporter lines with support from WS. ND carried out the yeast two hybrid experiments. ML provided reagents.

Visualization: AER, NM, MJP

Funding acquisition: AER, LS, ME, SH Project administration: AER, LS, ME, SH Data curation: AER

Supervision: AER, LS, ME, SH

Writing – original draft: AER, LS, ME, SH, NM, MP Writing – review & editing: AER, LS, ME, SH, NM, MP

## Data and materials availability

All original numerical, sequence and image data is included in the manuscript or freely available via the University of Edinburgh DataShare service (https://datashare.ed.ac.uk/handle/10283/3662 College of Science and Engineering/ School of Biological Sciences/ Institute of Molecular Plant Science/ The Plant Shape Lab). All code used for the sequence analysis is available on github: https://github.com/ThePlantShapeLab/Truffula. Seeds and plant material are available on request from Richardson (maize), Estelle (*P.patens*), Strader (Arabidopsis) and subject to MTAs.

## Supplementary Materials

### Materials and Methods

#### Maize plants

The *Truffula* seed was provided by Gerry Neuffer, and originally called Ts-2620, found in an EMS mutagenized Mo17 population. The phenotype of *Trf* was analyzed in backcrosses into W22 and Mo17 advanced to generation 10. Field grown plants were grown at UC Berkeley Gill Davis and UMass Amherst during summer field seasons. All plants used for molecular analysis were backcrossed into W22 10 times. Seedlings grown for RNA and DNA extractions were grown under greenhouse conditions in 30×45×2.5cm trays with peat free soils plus osmocote. Seedlings grown for immunolocalization and western blots were grown in temperature-controlled greenhouses with supplemental light in 30×45×2.5cm trays with peat-reduced osmocote compost with added fertilizer under 16/8 day/ night cycle.

#### DNA Extraction

Leaf discs were collected from 80 *Trf* and N siblings in the W22 backcross population (10 introgressions). High-quality gDNA was extracted using urea as described previously (*47*). In brief, genomic DNA was extracted using a urea-based extraction method which is as follows. Tissue was ground in urea extraction buffer (4M urea, 4M NaCl, 1M Tris pH8, 0.5M EDTA, 34mM n-lauroyl sarcosine). Then an equal volume of phenol:chloroform:isoamyl alcohol (25:24:1), was added and vortexed to mix. After centrifugation, the supernatant was decanted and mixed with an equal volume of chloroform. After centrifugation, the supernatant was decanted and mixed with 1/10^th^ volume of 4.4 M NH4OAc pH 5.2 and 0.7 volume of isopropanol. Strands of DNA were then collected and washed with 70% ethanol in a separate tube. The dried pellet was resuspended in 100ul of TE. DNA integrity was checked using gel electrophoresis and quantified using qubit before sending for library preparation and 150bp PE sequencing by Novogene. Sequencing was for 10 × coverage of the maize genome. DNA for SSR mapping and genotyping was extracted from 1.5cm of leaf tissue using the protocol described previously (*48*).

#### Whole-Genome Sequencing Bulk-segregant Analysis

Sequence data was downloaded via ftp from Novogene servers and passed through the following analysis pipeline: FASTPQ (*49*) > Bowtie2 (*50*) alignment to the B73_Reference_Nam5.0 genome > samtools mpileup variant calling (*51*) > SNPeff (*38*) and variant filtering in R, restricting to EMS type SNPs. Jupyter notebooks containing the code for WGS-BSA analysis is available at: https://github.com/ThePlantShapeLab

#### SSR Mapping

SSR markers were identified using the maize GDB database and used to genotype *Trf* phenotype plants from successive generations. (see Fig.2 for position, and Table S1 for primers). Custom SSR markers were identified and designed to refine the region further (see Table S1.) Genotyping PCRs were analyzed using metaphor agarose electrophoresis.

#### RNA Extraction

A segregating population of 3-week-old maize seedlings were first genotyped for *Trf* (see Table S1 for primers). The outer 2 leaves were removed and 0.5cm of the shoot apex was dissected and flash frozen in liquid nitrogen. Three replicate pools were created for *Trf* and N siblings, 10 individuals per pool. RNA was extracted from each pool using TRIzol® (Invitrogen) as described in the users manual. RNA quality was checked using gel electrophoresis and quantified using qubit before sending for library preparation and 150bp PE sequencing by Novogene.

#### RNAseq Analysis

Sequence data was downloaded via ftp from Novogene servers and passed through the following analysis pipeline: FASTPQ (*49*) > HiSAT2 (*52*) alignment to the *Zea mays* Reference NAM5.0 genome > FeatureCounts (*53*) > DeSeq2 (*54*) analysis of differential gene expression using genes with more than 5 counts in 3 or more samples. All analysis was carried out locally (3.6GHz 8-Core Intel Core i9 processor, 32GB RAM) using R packages combined with Jupyter notebooks. Jupyter notebooks containing the code for RNAseq analysis is available at: https://github.com/ThePlantShapeLab GO term analysis was carried out using the GAMER Maize annotations (*55*) and the GOSeq R package (*56*), significantly enriched GO terms had a Benjamini and Hochberg cutoff padj value of <0.05.

#### Antibody Production

Antibody was developed from the ZmARF28 full length coding sequence cloned into pDEST17 bacterial destination vector carrying the 6HIS tag for antigen production. Recombinant protein expression was carried out in *E. coli* Rosetta strain (BL21). After purification, a total of 200 µg of protein was sent to Cocalico Biologicals Inc. (Stevens, PA, USA) where two guinea pig immunizations were performed. To improve specificity of this antibody, purification from antiserum was carried out using a synthetic 20 amino acid peptide from position 505 to 524 which is a highly specific region to ZmARF28 (amino acids 505-524: RPFPNKISGTRSSTWVTADA) using magnetic beads from Invitrogen. A Cysteine (C) residue was added at the N-terminal end of the peptide to enable covalent binding to the beads used for antibody purification. To validate specificity of the purified antibody, a western blot was carried out following the described protocol in the next section, using 20 mg of soluble total protein from vegetative B73 maize shoot apical meristem.

#### Protein Extraction and Western Blots

3-week-old maize seedling shoot apices were harvested and flash frozen in liquid nitrogen, before grinding to a fine powder. 100mg of powdered tissue was then thawed in 1× SDS+DTT loading buffer and vortexed. Samples were heated at 95°C for 5 minutes, before loading on a 10% acrylamide gel for electrophoresis. Semi-dry transfer to a MeOH activated PVDF membrane in 1× Tris-Glycine/20% MeOH was carried out. Membranes were air dried, then reactivated in MeOH and washed in TBS/0.05% Tween for 5 minutes. Then, Blocked for 1 hour with 6% skimmed milk in TBS/0.05% Tween at room temperature. Membranes were incubated overnight in the primary AntiARF28 antibody diluted 1:250 in 1% milk/TBS/0.05% Tween. The following day membranes were washed (3× 5 mins in TBS/0.05% Tween) and incubated with 1:5000 Anti-guinea pig-HRP secondary antibody (Thermo Fisher Scientific catalog no. A18769) at room temperature with rocking for 2 hours. After washing 4× 5 mins in TBS/0.05% Tween membranes were incubated with Clarity Western ECL substrate (Biorad catalog no. 1705060) and exposed to x-ray film. Film was exposed to the membrane for 10 minutes before developing. This experiment was repeated four times with similar results.

#### Image Analysis and Figure Assembly

All images were analyzed in FIJI (*57*). Figures were assembled using Adobe Photoshop.

#### Statistical Analyses

All data were tested for normality using the Shapiro-Wilk test. Based on this analysis, data with a normal distribution was analyzed using a student’s T-test, or ANOVA where appropriate. Data that was not normally distributed was analyzed using the non-parametric Kruskal-Wallis test, followed by pairwise Wilcox tests. A Jupyter notebook containing the code for data analysis is available at: https://github.com/ThePlantShapeLab

#### Moss growth

Moss was grown on BCD minimal medium or BCD supplemented with either 5 mM ammonium tartrate (BCDAT) or 5 µM NAA (BCD+NAA) (*58*). BCDAT was used for routine growth. For growth assays, ∼1 mm^2^ pieces of tissue from chloronema-rich cultures of each strain were spotted at least twice in different positions on the plates containing all three media. The *nar* mutants were isolated in the *DR5:DsRed/NLS-4* background in the Gransden ecotype (*12*), and all subsequent analyses were carried out in either the Reute ecotype or a strain with the *DR5:DsRed* transgene introgressed from the semi-fertile Gransden strain into Reute through three crosses. During the course of this work, it was discovered that the *DR5:DsRed* expression is often silenced upon re-transformation, possibly due to the locus containing approximately 60 repeats of the transgene (*39*).

#### Moss mutant screen

The screen for *nar* mutants and mutant genome sequencing was described previously(*39*). Briefly, 30 plates of week-old lawns of *DR5:DsRed/NLS-4* on cellophane- overlain BCDAT media were irradiated with 150 mJ/cm^2^ of UV light in a Stratalinker UV Crosslinker (Stratagene). After 24 hours in darkness, the cellophanes were moved to new BCD+NAA plates, and the mutants were identified after 2–4 weeks. Confirmed mutants were screened for mutations in *Aux/IAA* and *DIAGEOTROPICA* genes by sequencing PCR products, and genomes of twelve of the strongest remaining mutants were sequenced. The full set of variants in the mutant genomes were filtered for those causing protein-altering changes in the same gene in two or more of the mutants. While both *narC29* and *narC18* both had the E263K substitutions in *ARFb4*, *narC18* also had a G608S substitution in class D ARF, *ARFd1* which likely accounts for the stronger phenotype. For simplicity, only *narC29* is shown.

#### CRISPR/Cas9

Plasmids for expressing one or two sgRNAs and Cas9 were assembled either as described in (*59*) or after modification to bypass the Gateway cloning step: the empty pENTR-L1L2-U6:BsaI insert was recombined into the pMH-Cas9-Gateway vector with LR Clonase II (Invitrogen), then the resulting plasmids backbone was replaced with that of pUK21 which lacks BsaI restriction sites and thus allows GoldenGate assembly of protospacers between the BsaI sites after the PpU6 promoter. A fragment of GFP was inserted between the BsaI sites to help distinguish recombinant clones from the parental vector. The *hph* selection gene was swapped for *nptII*, *aacC1*, or *ble* to allow selection with kanamycin/G418, gentamicin/G418, and Zeocin, respectively, in addition to Hyg. Protospacers were selected based on predicted specificity and efficiency using the http://crispor.tefor.net/ website (*60*). sgRNAs starting with G were preferred. Plasmids for making deletions contained dual *PpU6:sgRNA* genes targeting shortly after the start codons and before the stop codons. Oligo-mediated gene editing was achieved by designing double-stranded oligos matching the target but with the intended missense mutation(s) and silent mutations to create a restriction site and multiple mismatches in the PAM-proximal half of the sgRNA target. The oligos extended 21 bases before and after the mismatched bases.

#### Moss transformations

Protoplasts were prepared from week-old tissue and transformed as described (*58*) except that the post-heat-shock dilution was done by slowly adding 10 ml 8.5% mannitol all at once rather than 6.5 ml in 10 steps. Transformations included 15–25 µg of each plasmid and 250 pmol of annealed oligos. Deletion lines were identified by dramatically truncated PCR products and Sanger sequencing. Genome edited lines were identified by PCR across the edited site followed by digestion with the restriction enzyme whose site was added to the template oligos. PCR products with the expected digestion pattern were sequenced. In several instances, lines exhibiting the same mutant phenotype but lacking the restriction site change or lacking the phenotype but having the restriction site change were found to have incorporated either the missense mutation or the restriction site, respectively, without the other. In the first editing experiment to re-create the E266K and R269Q mutations in *PpARFb2*, it was discovered that the *proDR5:DsRed* transgene’s gentamicin selection cassette also confers resistance to the G418 selection agent and this precluded selection of the CRISPR plasmid. We instead screened the transformants for resistance to NAA. Some of the NAA-resistant lines had the intended mutation but many had small in-frame insertions and deletions in the region.

mYPet-tagging of endogenous loci was achieved by co-transforming a CRISPR plasmid targeting near the stop codon and a plasmid that included: 600-1500 bp homology arm of coding region preceding the stop codon fused in-frame to a linker (SRGGGGA) and *mYPet* (encoding a bright, monomeric yellow fluorescent protein), followed by the *Pisum rbcS* terminator, a *pro35S:Hyg:ter35S* selection cassette flanked by *loxP* sites, and a downstream homology arm consisting of the 3′ UTR and downstream sequences. Stable lines were identified by selecting for the CRISPR plasmid and the knock-in construct for 7 days, followed by 10–14 days of no selection, and 7 days of Hyg selection. Clean insertion lines were identified by 1) PCR amplification with primers matching before the upstream homology arm and downstream from the downstream homology arm with primers to *mYPet* and *ter35S* and 2) lack of PCR amplification across the insertion site nor with primers to Cas9. In the case of the *pparfb1^E266K^* and *pparfb2^E266K^*, the strength of the phenotypes increased upon integrating the *mYPet/Hyg* sequences. Removal of the *pro35S:Hyg:ter35S* selection cassette by *CRE/lox*-mediated excision reverted the phenotype to that prior to insertion suggesting that *pro35S* enhancers may act at a distance on the nearby promoters in moss as in Arabidopsis (*61*, *62*). All mYPet lines shown had their selection cassettes excised.

#### RT-PCR

Week-old *pparfb2^E266K^*, *ppiaa2^G325S^*, and the corresponding wild type (*DR5:DsRed*/3×Re) tissues were homogenized, plated on BCDAT plates overlain with cellophanes, and grown for six days. Eighteen hours before treatment, the cultures on cellophanes were moved to BCD plates to alleviate the antagonism on auxin response by ammonium. Approximately 100 mg of tissue were scraped from the cellophane and transferred in triplicate to wells of 6-well plates with BCD-agar media containing either 10 µM IAA or 0.01% ethanol with 1 ml liquid BCD media with the same concentrations of IAA and ethanol. After 1 hour, tissue was patted dry between paper towels and RNA was extracted using the *Quick*-RNA Plant Miniprep kit (Zymo Research) and a Mini-Beadbeater-16 (BioSpec Products) according to the instructions. 1.3 µg total RNA was reverse transcribed using the Maxima H– cDNA Synthesis Master Mix (ThermoFisher), diluted 8-fold with 0.1×TE, and 3 µl used per 20 µl SYBR-qPCR reaction using CFX Connect-96 Real-Time PCR Detection System (Bio-Rad).

#### Yeast Two Hybrid assay

Yeast interaction assays were carried out using the Matchmaker LexA Two-Hybrid System (Clontech). Interactions were indicated by β-galactosidase detection on SD/Gal/Raf/X-Gal plates. For testing BRD function, a C-terminally Myc-tagged full-length TPL2/Pp3c9_21250 cDNA was inserted into pGilda (LexA DBD) and full-length ARFb2 cDNA with and without the L614S and F615S substitutions were inserted into pB42AD.

#### Moss imaging

For growth assays, 3-week-old moss were photographed with a Nikon SMZ1500 and a DS-Ri1 camera. DsRed fluorescence was captured using a TRITC filter set. Confocal microscopy was performed using a Zeiss LSM880. Tissue was mounted in water or liquid BCD media, and images were captured for DIC, YFP fluorescence, and chlorophyll autofluorescence. FIJI was used for processing images and for measuring YFP signal (*57*). Nuclei were identified in the YFP or DIC channels, circled as freehand selections, and their integrated density measured. Note that not all nuclei in wild type—unlike the mutants—have detectable YFP signal and could not be measured, so their quantification is likely an overestimate and the difference between wild type and the mutants is an underestimate.

#### Cloning ARF2 and arf2T98N constructs

DNA encoding 4 GS repeats, a porcine teschovirus-1 2A (P2A) cleavage peptide (*6*), and mScarlet3 were synthesized using TWIST biosciences fragment synthesis. The P2A- mScarlet3 insert was amplified by PCR using Q5 2× master mix (NEB). The protoplast expression vector pLCS107 was linearized by PCR using Q5 2× master mix (NEB). The insert was cloned into linearized pLCS107 using NEBuilder Hifi cloning reactions to generate pUBQ-mNEON-P2A-mScarlet. The arf2T298N variant mutation in the ARF2 CDS in pENTR ARF2 by PCR amplification and infusion cloning (Takara BioSciences). Both wild-type and T298N ARF2 were amplified from a pENTR backbone by PCR and cloned in-frame with mNEON and the GS linker using NEBuilder Hifi cloning. The pUBQ- mNEON-2A-mScarlet backbone was linearized by PCR.

#### Arabidopsis Protoplast Expression and Quantification

Arabidopsis mesophyll protoplasts were isolated from 14-day-old Col-0 leaves. 100,000 cells were transformed with 20-30 μg of plasmid DNA carrying mNEON-ARF2-P2A- mScarlet3 or mNEON-arf2T298N-P2A-mScarlet3 constructs using the tape-sandwich method and incubated for 16 hrs in the dark (*63*). Transformed cell populations were scored using the Beckman Coulter Cytoflex S Flow Cytometer and CytExpert software. A back gating strategy was taken to identify the population of intact protoplasts. Cells expressing mScarlet3 reporters were first identified by comparing transformed mNEON- ARF2-P2A-mScarlet3 cells to untransformed cells using the using the Y610 channel (Ex: 561 nm Em: 610±20 nm, 1000 gain) and then back gated on FSCvSSC. Cells were treated with 100µM cycloheximide diluted in protoplast media. for the indicated times. A minimum of 162 mScarlet3 positive cells from six independent transformations were used to collect mScarlet3 reporter levels. The mNEON levels were collected using the B525 channel (Ex: 488 nm Em: 525±40 nm, 69 gain). FCS files for mScarlet3 positive cell populations were generated by Cytoflex and analyzed using FlowKit (*64*), NumPy (*65*), and Pandas(*66*) (https://zenodo.org/record/7979740) packages in python with custom scripts. Ratios were determined by taking the mNEON signal divided by the mScarlet3 signal on a per cell basis. The mean ratio of each replicate at timepoint 0 was generated and all cells from that replicate were normalized to this value. A minimum of 2166 cells across all 6 replicates were collected. Graphs were generated using seaborn (*67*) and matplotlib (*68*) in python.

#### ARF DNA-binding Domain Structural Predictions

Protein structure predictions of ARF DNA-binding domains as defined in Boer et al 2014 (*17*) from wild-type and mutant ZmARF28, PpARFb2, and PpARFb4 were generated using CollabFold v1.5.2 (https://colab.research.google.com/github/sokrypton/ColabFold /blob/main/AlphaFold2.ipynb) (*69*) accessed on Aug 8 2023 using default conditions. The top model from each prediction was visualized with PyMol 2.5 (http://www.pymol.org).

#### Phylogenetic tree

Protein sequences for class B ARFs from 13 species were identified by reciprocal BLAST searches and aligned using T-COFFEE (v11.00.8cbe486) (*70*). Non-conserved regions were trimmed using Mesquite (v3.70) (*71*), and a tree was inferred using MrBayes (v3.2.7) (*72*) with settings aamodelpr=mixed, rates=invgamma, 2 runs of 4 chains for 1000000 generations.

Databases: *Alsophila spinulosa* (*73*); *Amborella trichopoda* (*74*); *Arabidopsis thaliana* (*75*); *Ceratodon purpureus* (*76*); *Ceratopteris richardii* (*77*); *Diphasiastrum complanatum*(*78*); *Marchantia polymorpha* (*79*); *Physcomitrium patens* (*80*); *Selaginella moellendorffii* (*81*); *Solanum lycopersicum* (*82*); *Takakia lepidozioides* (*83*); *Thuja plicata* (*84*) ; *Zea mays* (*85*). Class B ARF sequences were not found in the three published genomes nor in the 1KP transcriptomes for hornwort species (*86–88*).

## Supplementary Materials

**Fig. S1.**
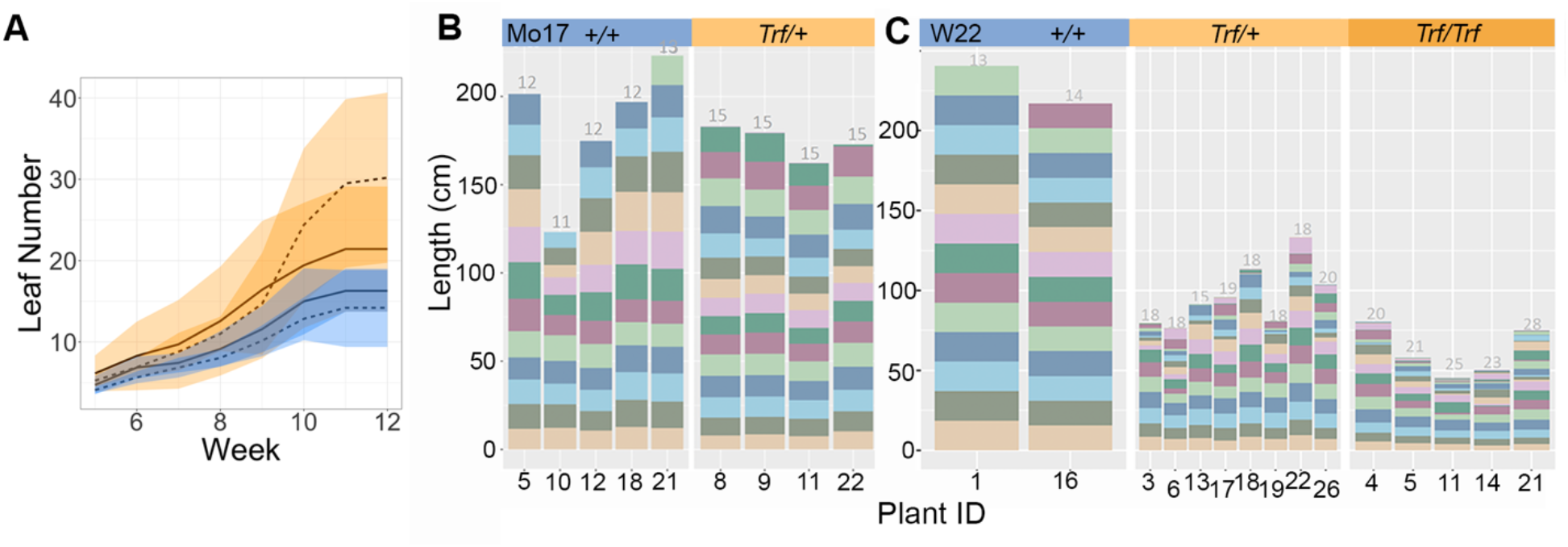
*Truffula* mutants have defects in leaf initiation and internode elongation (**A**) Leaves are initiated at a faster rate in heterozygous *Trf* mutants (orange) compared to normal siblings (blue) in both Mo17 (solid line) and W22 (dashed line) backgrounds. Line: mean leaf number, color shading: upper and lower bounds (mean ± 1.5 sd). *Trf* mutants (orange panels) in both Mo17 (**B**) and W22 (**C**) backgrounds have defects in internode elongation, with internode size shorter and more variable than normal siblings. Total bar height: plant height, color bars: internode length, numbers above the columns: number of visible internodes in each mature plant.

**Fig. S2.**
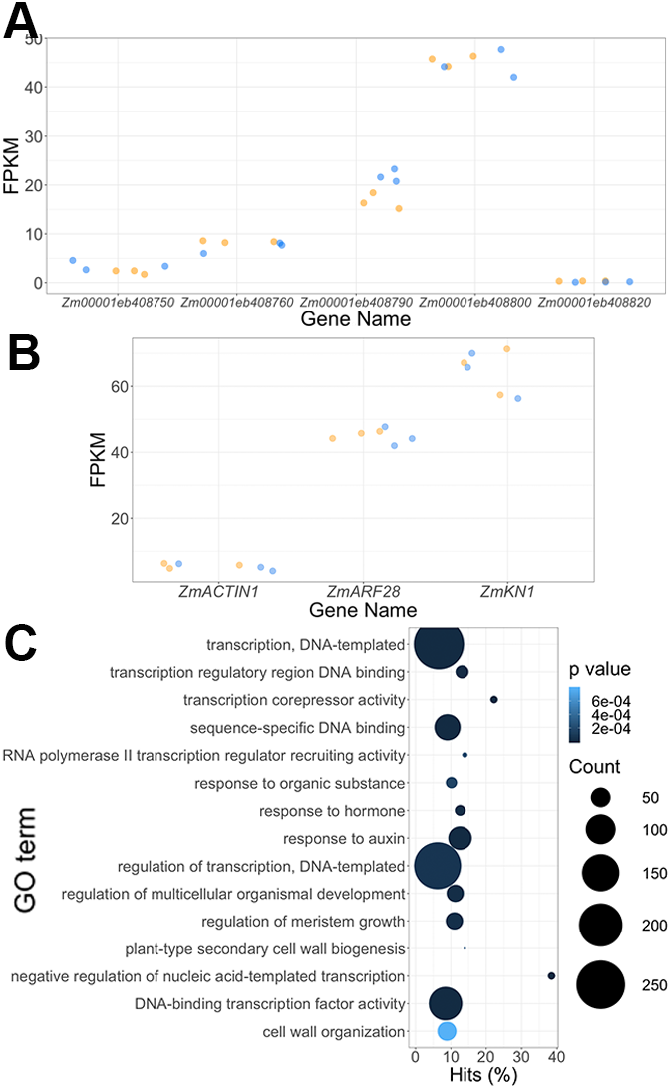
*Truffula* mutants do not show changes in ZmARF28 expression, and have defects in auxin signalling transcriptional responses (**A-B**) FPKM plots of specific genes in an RNAseq experiment of the vegetative shoot apices of heterozygous *Truffula* (*Trf/+,* orange) and normal (*+/+*, blue) siblings. (**A**) Expression of genes in the interval that have >5 counts in >=2 samples. (**B**) Expression of *ZmACTIN1* (Zm00001eb348450), *ZmARF28* (Zm00001e408800), and *ZmKNOTTED1* (Zm00001eb055920) (**C**) GO term enrichment analysis of significantly differentially expressed genes (padj<0.05, log2FoldChange >1, <-1) between *Trf/+* and *+/+* vegetative shoot apices.

**Fig. S3.**
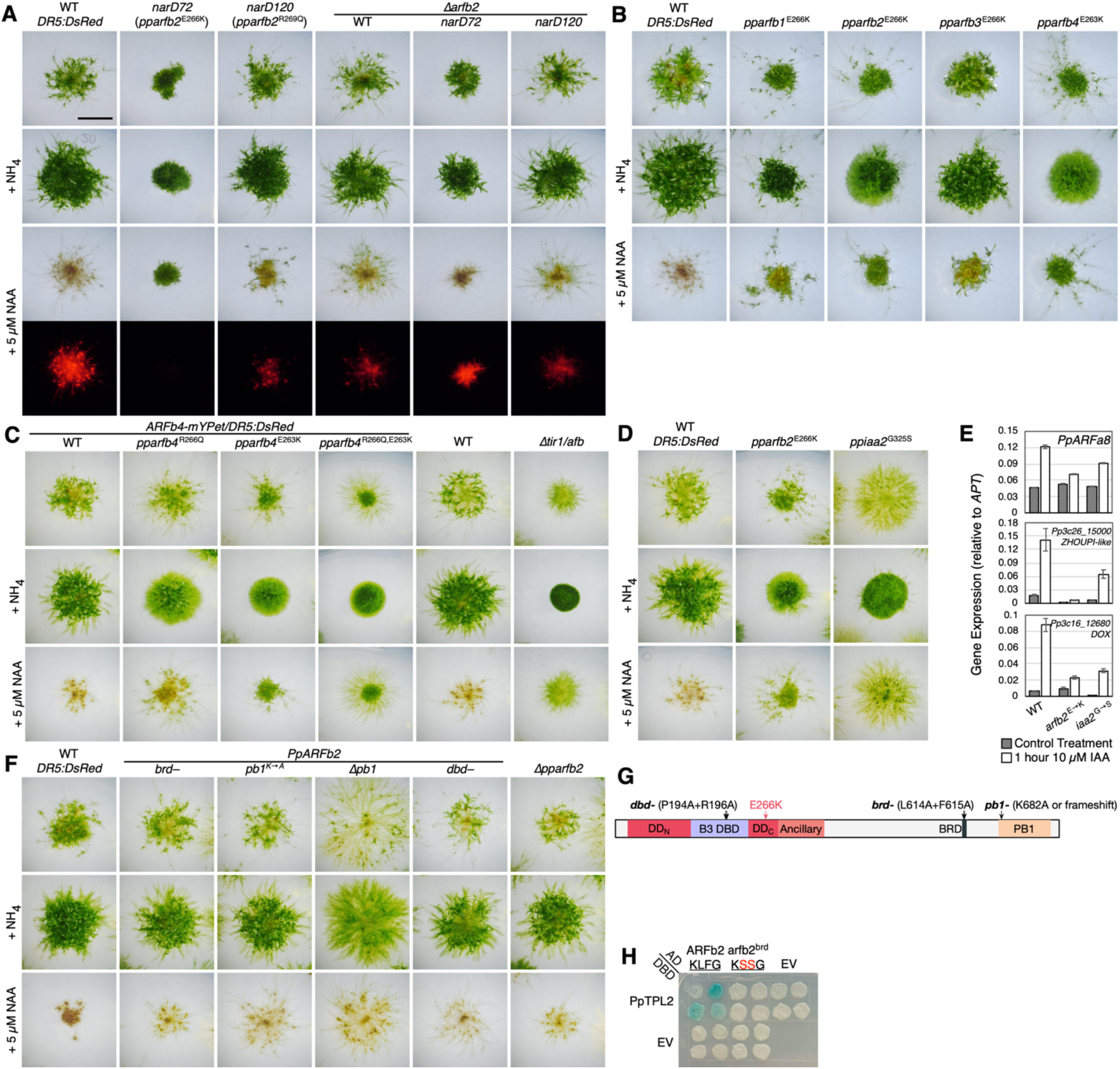
Mutations in *PpARFb* genes cause auxin resistance (**A**) Comparison of wild type, *nar* mutants, and *Δpparfb2* deletion mutants after 21 days of growth on standard medium (BCD) and BCD supplemented with 5 mM ammonium tartrate or 5 µM NAA. The plants grown on NAA were also imaged for RFP fluorescence from the *proDR5:DsRed* auxin-response reporter. Note that the leafy shoots in *narD120* are no longer present after deleting the mutant *pparfb2* gene. Scale bar is 5 mm. (**B**) Phenotypes of 21-day-old wild type and gene-edited mutants with E-to-K substitutions in all four class B ARFs showing leafy shoots on NAA media. (**C**) 21-day phenotypes as in panel (A) of the endogenously mYPet-tagged *PpARFb4* locus with either the R266Q, E263K, and both together introduced by gene editing. The *Δtir1/afb* quadruple mutant is shown for comparison. (**D–E**) Auxin-induced gene expression in wild type, *pparfb2^E266K^*, and *ppiaa2^G325S^* (a partially stabilized Aux/IAA degron allele). (**D**) 21-day phenotypes of the strains used in gene expression assay. (**E**) Gene expression of the class-A ARF *PpARFa8* gene, a *ZHOUPI-like bHLH* gene, and a member of the iron/ascorbate-dependent oxidoreductase family. (**F-G**) (**F**) Mutations affecting BRD, PB1, and DBD functions of PpARFb2 have little effect on auxin response, but those with the frameshift mutation at the start of the PB1-encoding region exhibit an increase in protonemal spreading which may indicate auxin hypersensitivity. (**G**) Diagram illustrating the positions of the mutations in panel (**F**) and Fig 3D. (**H**) Yeast-two-hybrid assay showing that wild-type PpARFb2 interacts with TOPLESS homolog PpTPL2 and that this interaction is abolished in the L614S+F615S *brd-* mutant

**Fig. S4.**
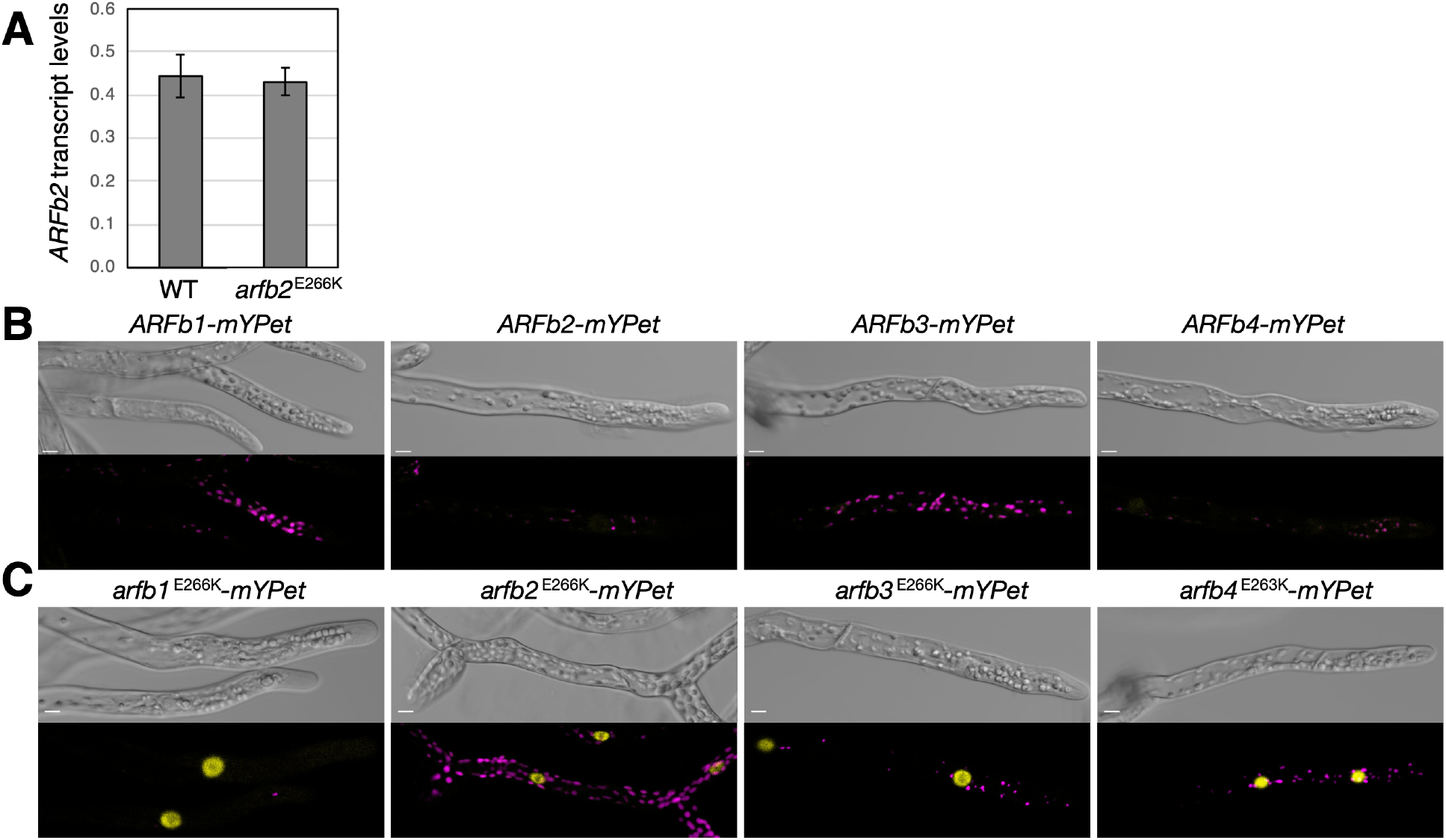
Substitutions in *PpARFb* increase protein stability (**A**) *PpARFb2* transcript levels are not significantly altered by the E266K mutation in tissue grown for 1 week on BCDAT medium. Transcript levels are normalized to that of the *APT* gene. (**B–C**) YFP fluorescence from wild-type and gene-edited mYPet-tagged *PpARFb* loci. DIC images and a merged micrograph for the YFP channel (yellow) and chlorophyll autofluorescence (magenta). The 514 nm excitation laser power was reduced four-fold for acquiring YFP signal from gene-edited lines in panel (C) compared to wild-type loci in panel (B). Scale bars are 10 µm. (**B**) YFP signal is very low or absent for tagged wild-type PpARFb-mYPet proteins. (**C**) E-to-K substitutions in all four PpARFb-mYPet lines increased YFP fluorescence dramatically.

**Fig. S5.**
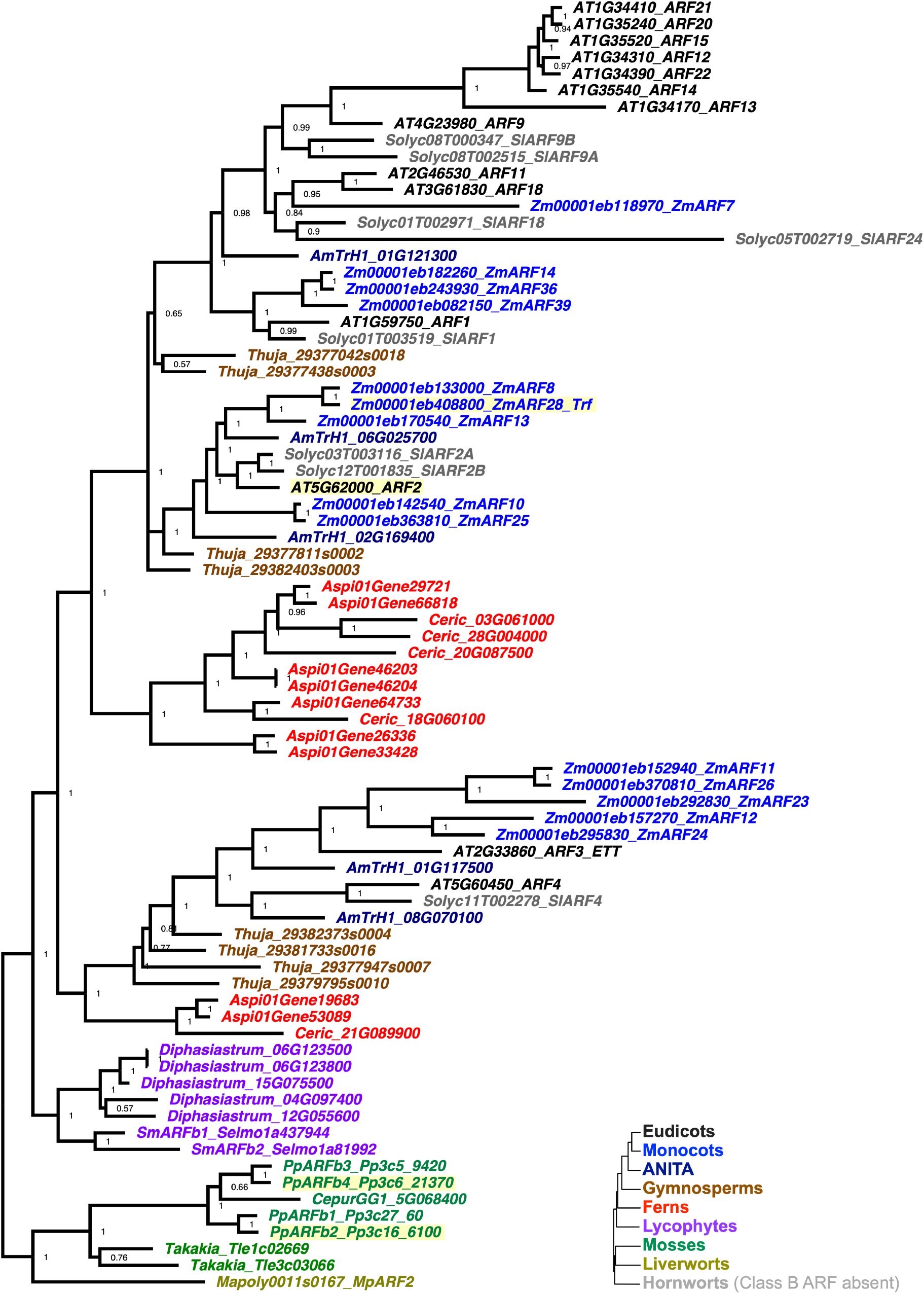
Bayesian-inferred phylogenetic tree of class B ARFs from thirteen land plant species Proteins are colored according to plant clades illustrated in the inset tree showing their relationships. The ZmARF28, AtARF2, PpARFb2, and PpARFb4 sequences are highlighted in yellow. Node labels indicate posterior probabilities supporting the enclosed clades.

**Table S1.**
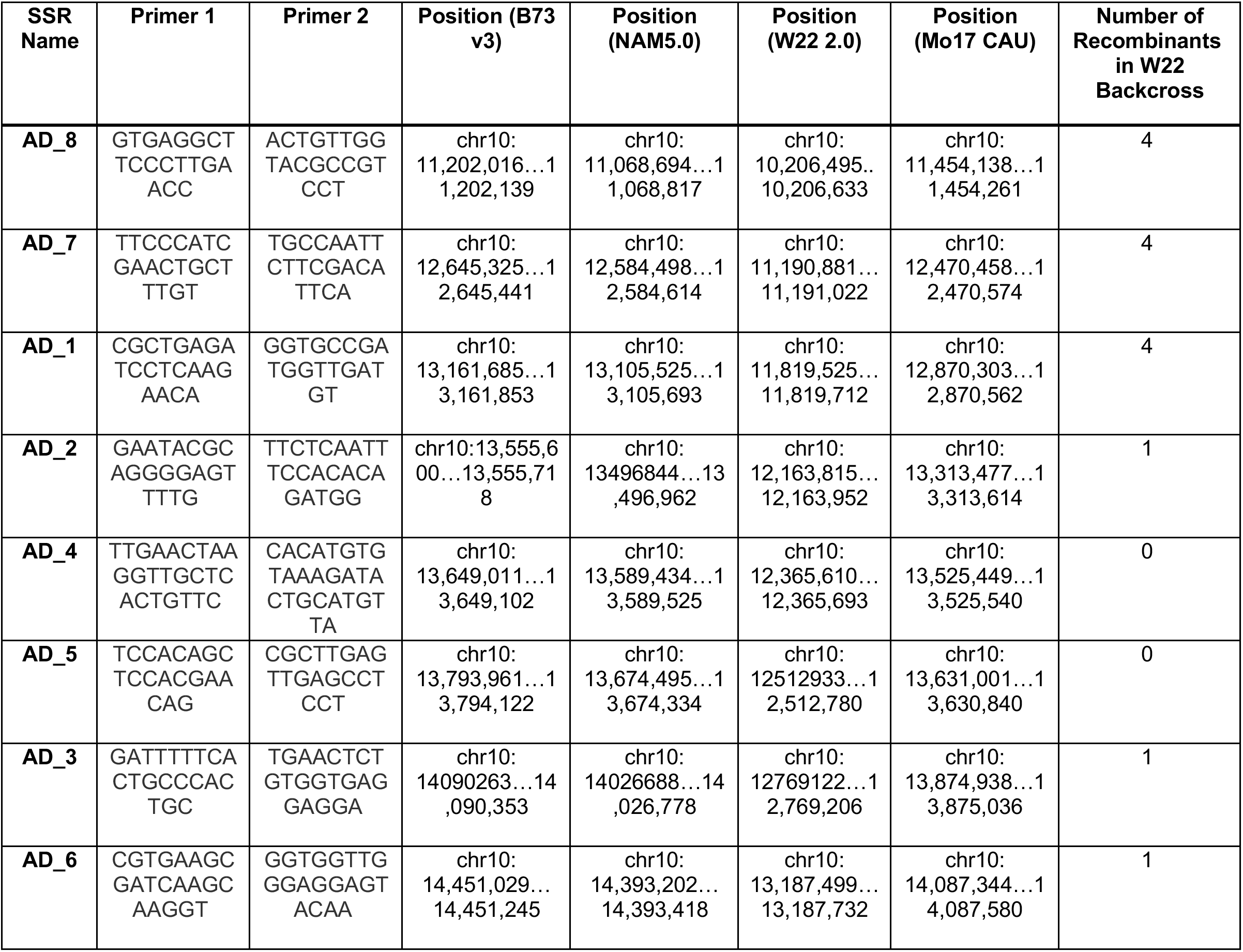
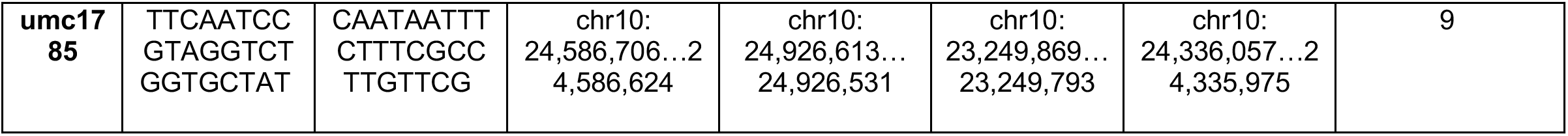
SSR Primers used in mapping of the *Trf* mutation. Reference genomes used: Zm-B73-REFERENCE-NAM-5.0, Mo17 CAU Assembly (Zm00014a), B73 RefGen_v3 (MGSC), W22 NRGene 2.0 assemblies

**Table S2.**
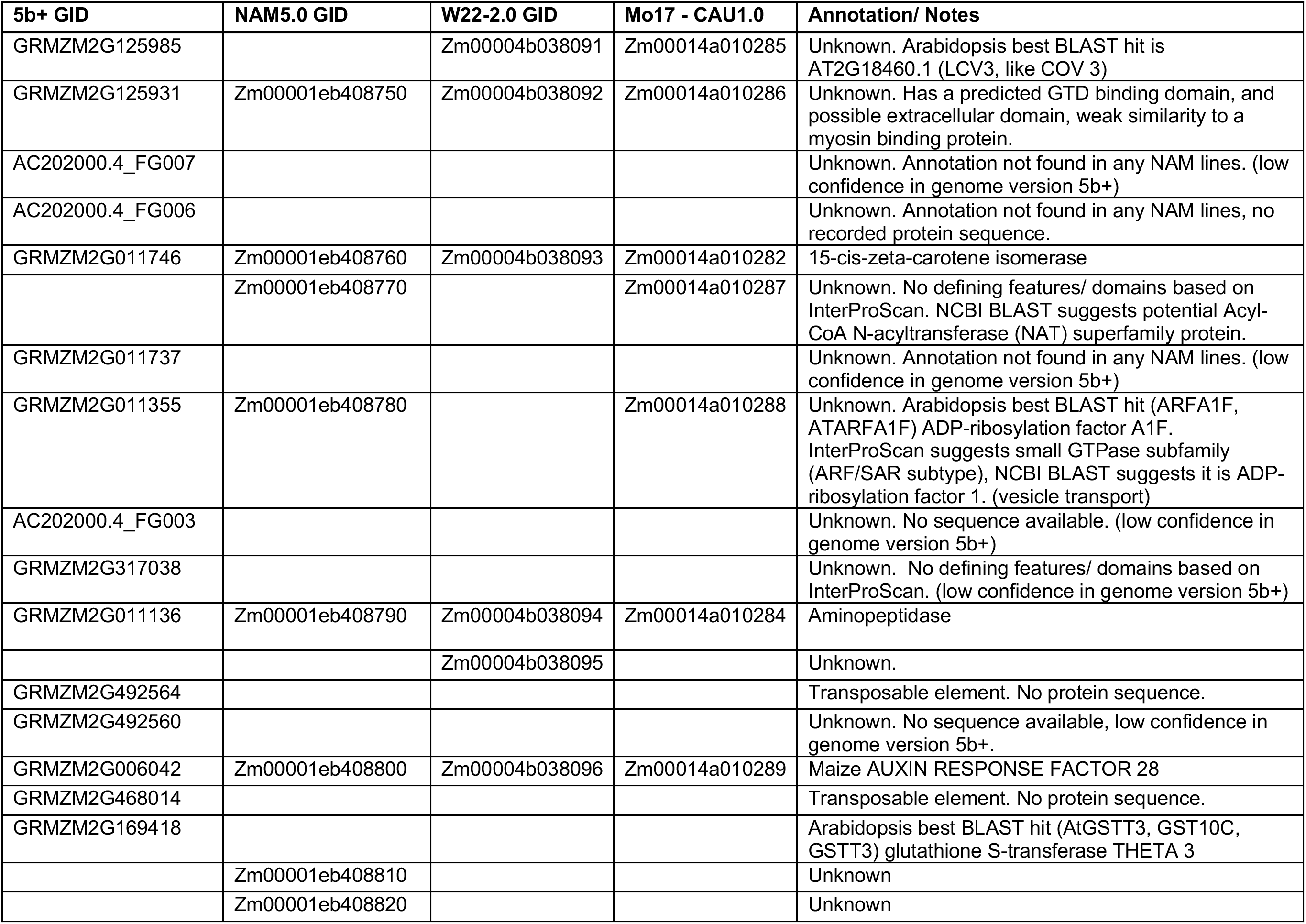
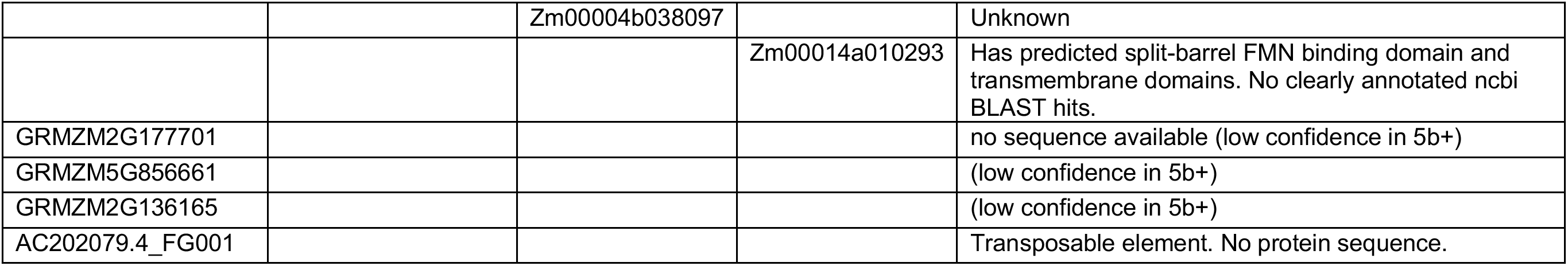
Genes annotated in the *Trf* mapping interval as determined by SSR primer mapping in the 5b+, NAM5.0, W22-2.0, and Mo17-CAU1.0 reference genomes available on maize GDB. Up to date for March 2023 versions of each genome annotation. See Table S1 for precise mapping interval location for each genome.

| 5b+ GID | NAM5.0 GID | W22-2.0 GID | Mo17 - CAU1.0 | Annotation/ Notes |
| --- | --- | --- | --- | --- |
| GRMZM2G125985 |  | Zm00004b038091 | Zm00014a010285 | Unknown. Arabidopsis best BLAST hit is AT2G18460.1 (LCV3, like COV 3) |
| GRMZM2G125931 | Zm00001eb408750 | Zm00004b038092 | Zm00014a010286 | Unknown. Has a predicted GTD binding domain, and possible extracellular domain, weak similarity to a myosin binding protein. |
| AC202000.4_FG007 |  |  |  | Unknown. Annotation not found in any NAM lines. (low confidence in genome version 5b+) |
| AC202000.4_FG006 |  |  |  | Unknown. Annotation not found in any NAM lines, no recorded protein sequence. |
| GRMZM2G011746 | Zm00001eb408760 | Zm00004b038093 | Zm00014a010282 | 15-cis-zeta-carotene isomerase |
|  | Zm00001eb408770 |  | Zm00014a010287 | Unknown. No defining features/ domains based on InterProScan. NCBI BLAST suggests potential Acyl-CoA N-acyltransferase (NAT) superfamily protein. |
| GRMZM2G011737 |  |  |  | Unknown. Annotation not found in any NAM lines. (low confidence in genome version 5b+) |
| GRMZM2G011355 | Zm00001eb408780 |  | Zm00014a010288 | Unknown. Arabidopsis best BLAST hit (ARFA1F, ATARFA1F) ADP-ribosylation factor A1F. InterProScan suggests small GTPase subfamily (ARF/SAR subtype), NCBI BLAST suggests it is ADP-ribosylation factor 1. (vesicle transport) |
| AC202000.4_FG003 |  |  |  | Unknown. No sequence available. (low confidence in genome version 5b+) |
| GRMZM2G317038 |  |  |  | Unknown. No defining features/ domains based on InterProScan. (low confidence in genome version 5b+) |
| GRMZM2G011136 | Zm00001eb408790 | Zm00004b038094 | Zm00014a010284 | Aminopeptidase |
|  |  | Zm00004b038095 |  | Unknown. |
| GRMZM2G492564 |  |  |  | Transposable element. No protein sequence. |
| GRMZM2G492560 |  |  |  | Unknown. No sequence available, low confidence in genome version 5b+. |
| GRMZM2G006042 | Zm00001eb408800 | Zm00004b038096 | Zm00014a010289 | Maize AUXIN RESPONSE FACTOR 28 |
| GRMZM2G468014 |  |  |  | Transposable element. No protein sequence. |
| GRMZM2G169418 |  |  |  | Arabidopsis best BLAST hit (AtGSTT3, GST10C, GSTT3) glutathione S-transferase THETA 3 |
|  | Zm00001eb408810 |  |  | Unknown |
|  | Zm00001eb408820 |  |  | Unknown |
|  |  | Zm00004b038097 |  | Unknown |
|  |  |  | Zm00014a010293 | Has predicted split-barrel FMN binding domain and transmembrane domains. No clearly annotated ncbi BLAST hits. |
| GRMZM2G177701 |  |  |  | no sequence available (low confidence in 5b+) |
| GRMZM5G856661 |  |  |  | (low confidence in 5b+) |
| GRMZM2G136165 |  |  |  | (low confidence in 5b+) |
| AC202079.4_FG001 |  |  |  | Transposable element. No protein sequence. |

**Table S3.** Significant and moderate effect SNP variants identified in the *Trf* mapping interval using SnpEff, based on the NAM5.0 maize genome annotation.

| **NAM5.0 Gene ID** | **SNP Variant** | **Predicted amino acid change** | **Notes** |
| --- | --- | --- | --- |
| Zm00001eb408750 | chr10:<br>13,501,412<br>C>T | Arg331Lys in T001 | natural variant in Mo17 |
| Zm00001eb408750 | chr10:<br>13,501,725<br>G>T | Leu227Met in T001 | natural variant in Mo17 |
| Zm00001eb408750 | chr10:<br>13,501,946<br>C>T | Cys153Tyr in T001 | natural variant in Mo17 |
| Zm00001eb408750 | chr10:<br>13,503,214<br>C>A | Ala62Ser in T001 | natural variant in Mo17 and W22 |
| Zm00001eb408800 | chr10:<br>13,676,774<br>G>A | T004 Ser281Asn, T005 Ser280Asn, T002 Ser281Asn, T006 Ser281Asn, T001 Ser281Asn | conserved serine across B73, W22, Mo17 |
| Zm00001eb408800 | chr10:<br>13,678,064<br>G>A | T003 Thre558Ile, T005 Thr558Ile, T002 Thr559Ile, T004 Thr559Ile, T006 Thr559Ile, T001 Thr559Ile | Variable residue across B73, W22, Mo17 |
| Zm00001eb408820 | chr10:<br>13,953,240<br>C>A | T001 Asp99Glu | Gene annotation absent from Mo17 and W22 reference genomes, CML52 has Glu at this residue. |

Table S4. All significantly differentially expressed genes (padj<0.05) in the RNAseq comparison of *Trf* vs normal sibling vegetative shoot apices.

(see excel spreadsheet)

**Table S5.**
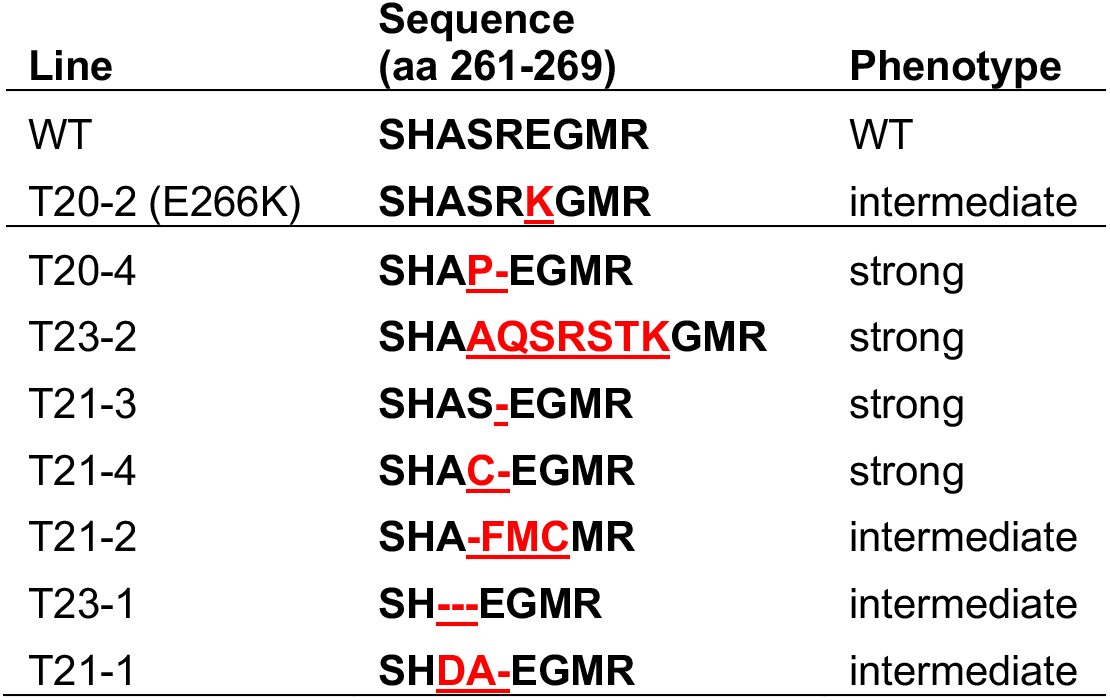
In-frame deletions and insertions (red) between amino acids 261-269 in *PpARFb2* confer either intermediate phenotypes similar to *pparfb2*^E266K^ or strong phenotypes similar to *pparfb2*^E266K+R269Q^ in *Physcomitrium patens*.

**Table S6.**
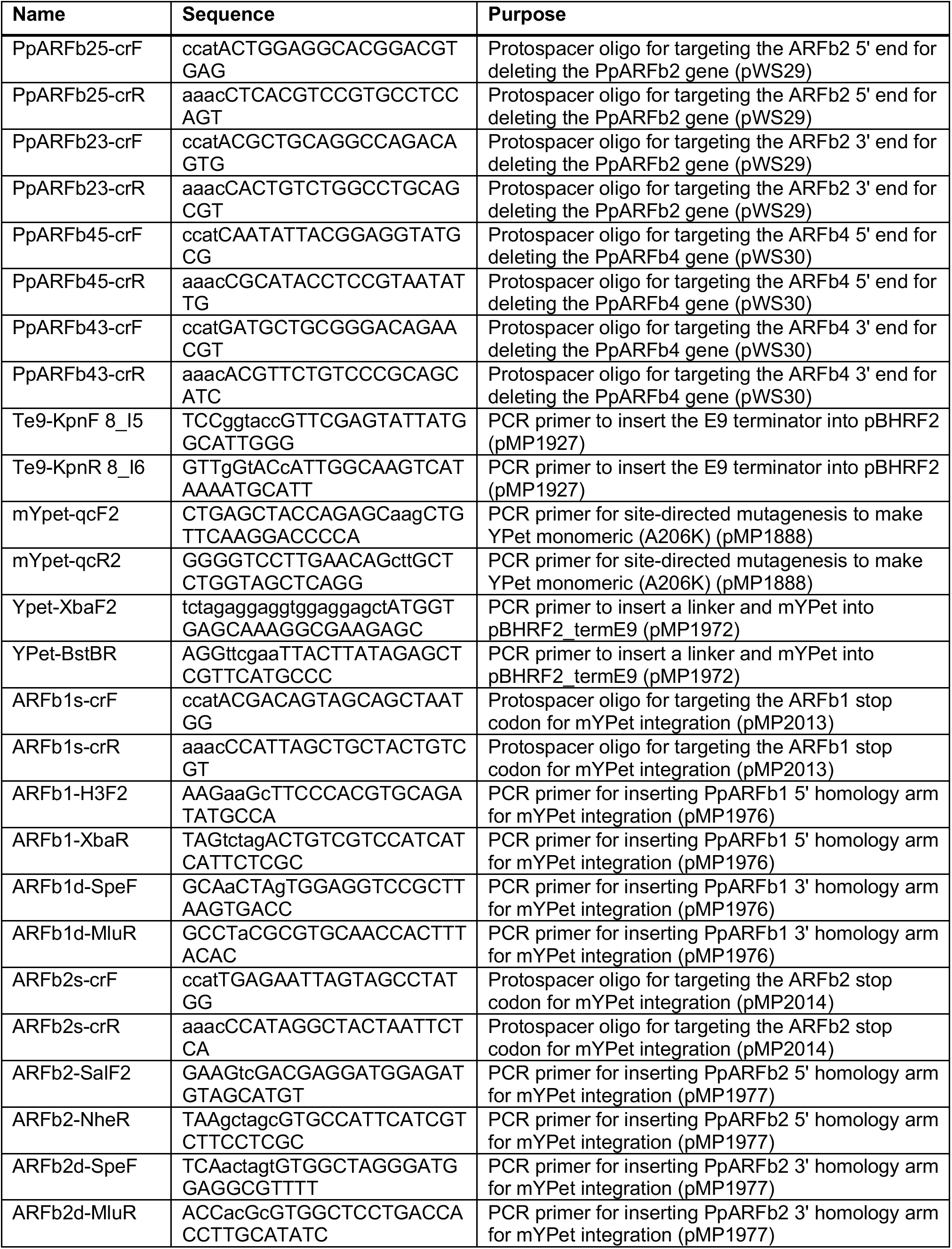
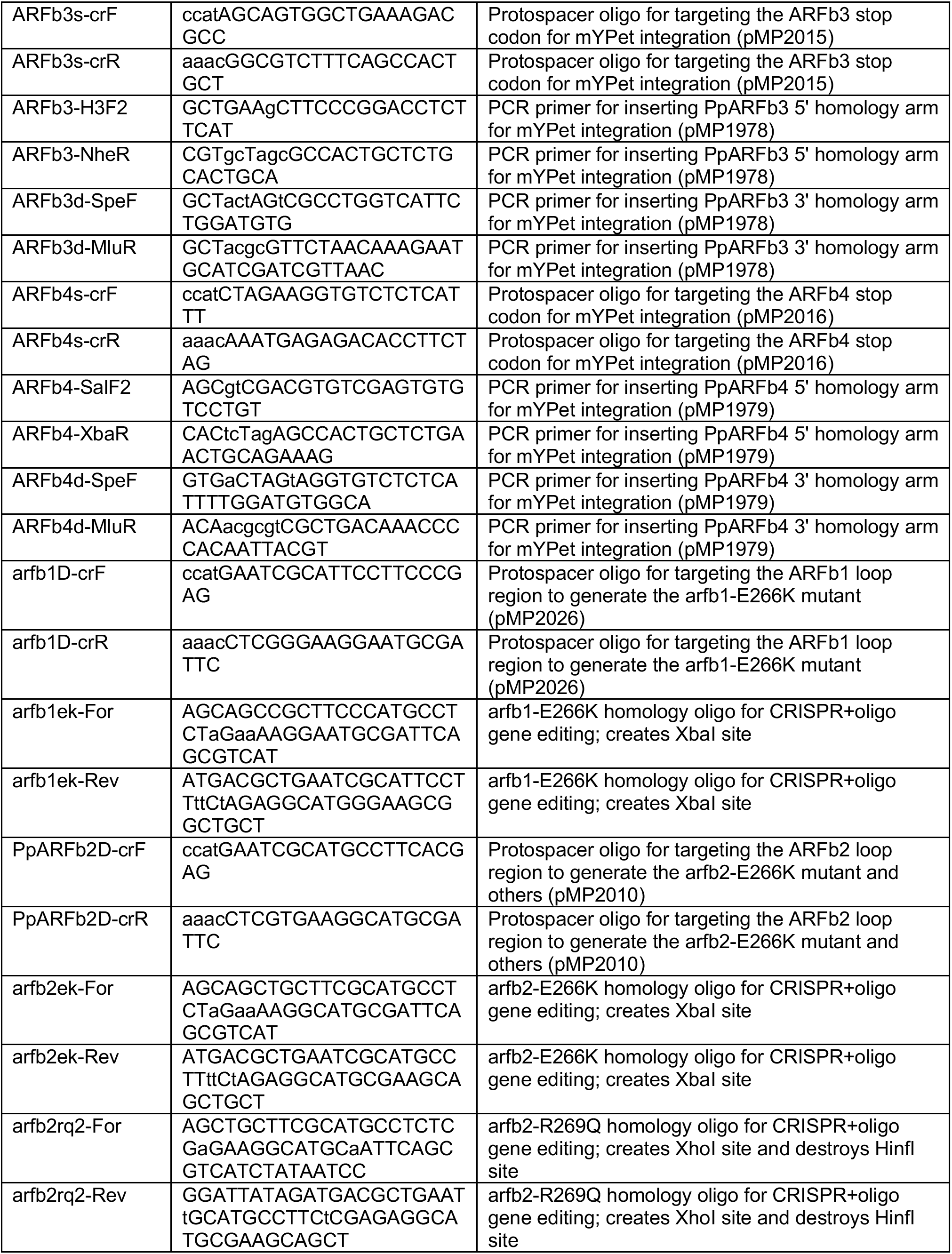
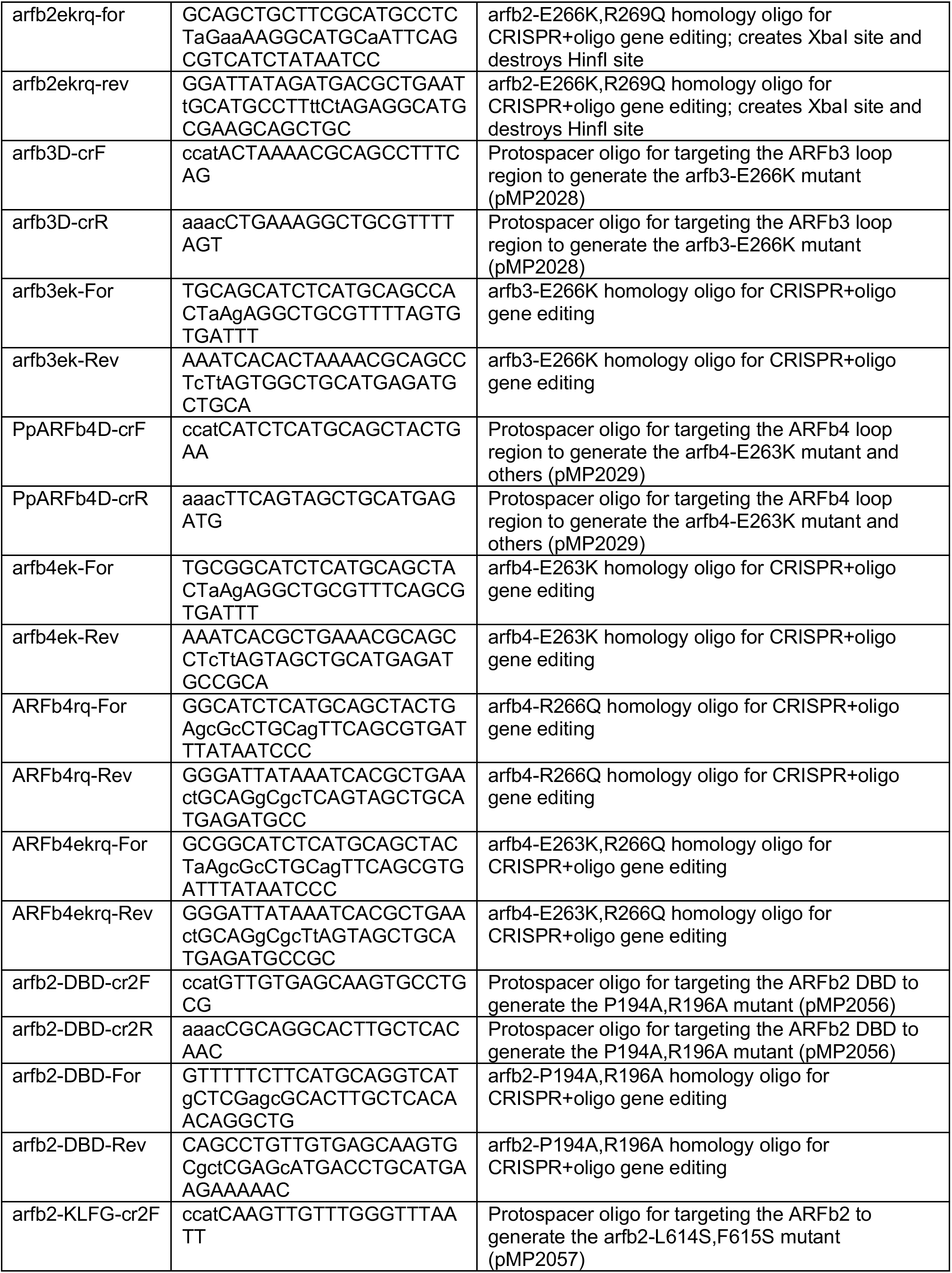
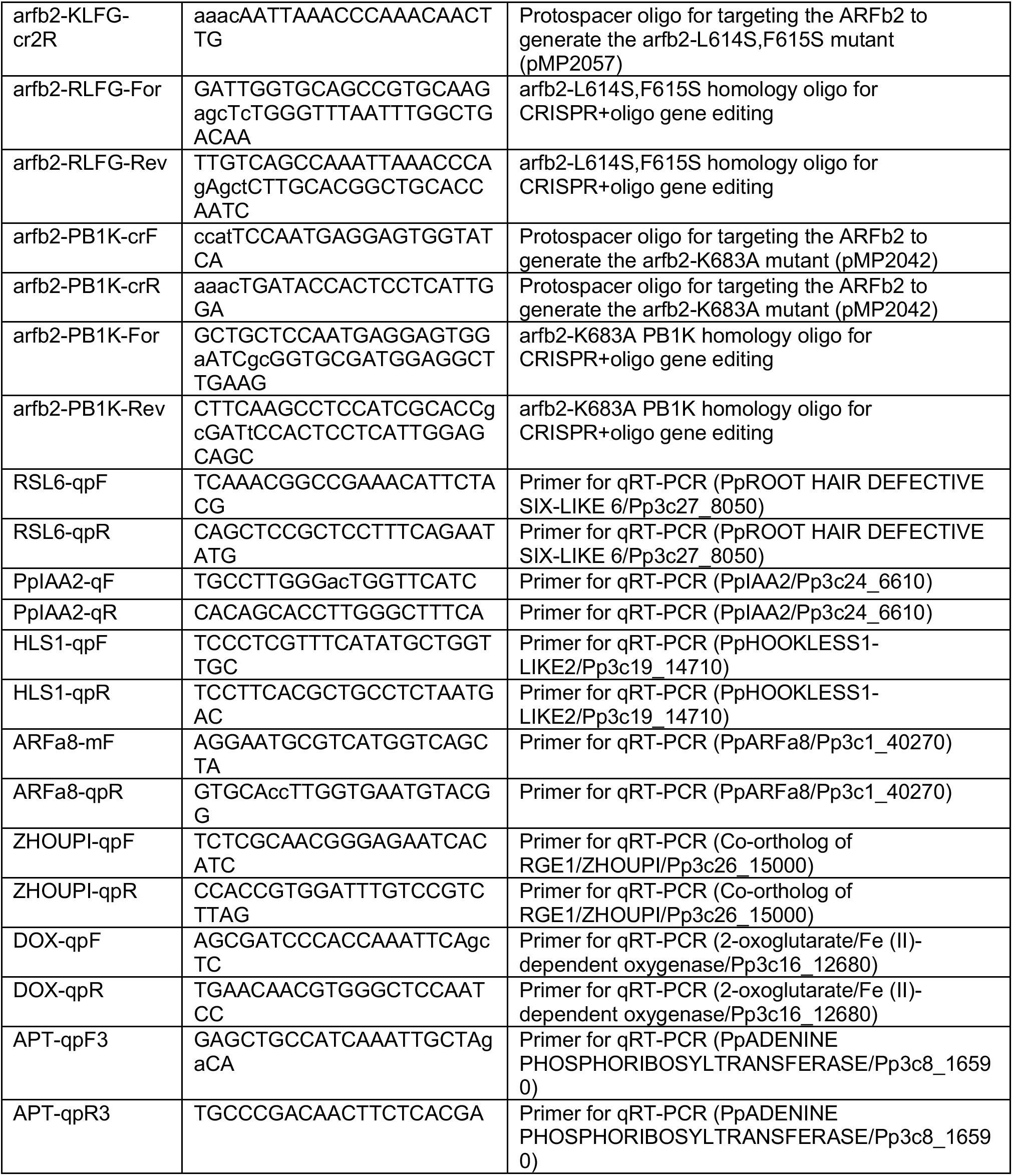
Primers and oligos used in the *Physcomitrium patens* work.

